# Lack of the IFN-γ signal leads to lethal *Orientia tsutsugamushi* infection in mice with skin eschar lesions

**DOI:** 10.1101/2024.02.05.578851

**Authors:** Yuejin Liang, Hui Wang, Keer Sun, Jiaren Sun, Lynn Soong

## Abstract

Scrub typhus is an acute febrile disease due to *Orientia tsutsugamushi* (*Ot*) infection and can be life-threatening with organ failure, hemorrhage, and fatality. Yet, little is known as to how the host reacts to *Ot* bacteria at early stages of infection; no reports have addressed the functional roles of type I versus type II interferon (IFN) responses in scrub typhus. In this study, we used comprehensive intradermal (i.d.) inoculation models and two clinically predominant *Ot* strains (Karp and Gilliam) to uncover early immune events. Karp infection induced sequential expression of *Ifnb* and *Ifng* in inflamed skin and draining lymph nodes at days 1 and 3 post-infection. Using double *Ifnar1*^-/-^*Ifngr1*^-/-^ and *Stat1*^-/-^ mice, we found that deficiency in IFN/STAT1 signaling resulted in lethal infection with profound pathology and skin eschar lesions, that resembled to human scrub typhus. Further analyses demonstrated that deficiency in IFN-γ, but not IFN-I, resulted in impaired NK cell and macrophage activation and uncontrolled bacterial growth and dissemination, leading to metabolic dysregulation, excessive inflammatory cell infiltration, and exacerbated tissue damage. NK cells were found to be the major cellular source of early IFN-γ, contributing to the initial *Ot* control. In vitro studies with dendritic cell cultures revealed a superior antibacterial effect offered by IFN-γ than IFN-β. Comparative in vivo studies with Karp- and Gilliam-infection revealed a crucial role of IFN-γ signaling in protection against progression of eschar lesions and *Ot* infection lethality. Additionally, our i.d. mouse models of lethal infection with eschar lesions are promising tools for immunological study and vaccine development for scrub typhus.

**Summary:** Scrub typhus can lead to severe complications and even fatality if not treated properly; however, the early host immune responses to *Ot* bacterium infection remain unclear. This study focused on the functional roles of IFNs in i.d. inoculation mouse models of scrub typhus. We found that mice lacking IFN receptors were highly susceptible to *Ot* infection, which resulted in severe pathology and skin eschar lesions that resembled to human scrub typhus. Further investigation revealed that the lack of IFN-γ, but not IFN-I, resulted in dysregulated innate immune responses, leading to uncontrolled bacterial burdens and tissue damage. Using IFN-γ reporter mice and neutralizing antibody treatment, we confirmed that NK cells were the major source of early IFN-γ, and thus played a key role in controlling *Ot* dissemination. Moreover, our comparative studies with two *Ot* strains revealed bacterium strain- and dose-dependent eschar formation and disease severity. In conclusion, our study highlights the crucial role of IFN-γ signaling in ensuring host protection against *Ot* infection. Our mouse models resemble skin eschar lesions and lethal infections observed in human disease, offering potential for future immunological studies on scrub typhus.

## Introduction

Scrub typhus is an infectious disease caused by the obligately intracellular bacterium, *Orientia tsutsugamushi* (*Ot*). Transmitted through the bite of a larval *Leptotrombidium* mite (commonly known as a chigger), this bacterium infects approximately 1 million people per year in the Asia-Pacific region, which houses over one-third of the world’s population and is referred to as the “tsutsugamushi triangle”[1, 2]. Typical symptoms among scrub typhus patients include skin eschar (a necrotic lesion with a black crust and an erythematous halo), regional lymphadenopathy, fever, and flu-like symptoms[2, 3]. *Ot* can spread to nearly all organs, leading to hyperinflammation, tissue damage, and multi-organ failure, with a median mortality rate of approximately 6% to 9% in untreated cases[3, 4]. Recent studies have identified scrub typhus in areas previously believed to be free of this disease, including South America and Africa[5, 6]. In 2023, the molecular detection of *Orientia* species was reported, for the first time, in free-living *Eutrombicula* chiggers collected in North Carolina[7], which highlights the concerning potential of human illness related to *Ot* infection in the United States. At present, no effective vaccines are available for scrub typhus, partly due to the antigenic diversity among various *Ot* strains and undefined roles of host immune responses in controlling or promoting the infection[8, 9].

In pursuit of a better understanding of host responses to *Ot* infection, we have recently developed several mouse models to investigate antibacterial immunity and immunopathogenesis. For example, the intravenous (i.v.) infection route results in the wide-spread infection of endothelial cells, leading to a lethal infection in both inbred C57BL/6j (B6) and outbred CD-1 mice, with pathological and immunological features resembling those observed in human scrub typhus[10–13]. However, the i.v. mouse model does not recapitulate the natural route of infection initiated through skin inoculation and is unsuitable for studying innate immunity and bacterial dissemination at early disease stages. To address such limitations, our research group and others established different intradermal (i.d.) mouse models (via inoculation in the ear or footpad), which helped understand infection characteristics relevant to human scrub typhus, such as the emergence of regional lymphadenopathy, innate immune cell activation, and the systemic dissemination of bacteria to distant organs[14–16]. The ear i.d. model has allowed us to uncover that dermal dendritic cells (DCs) and macrophages (MФs) can uptake *Ot* bacteria for dissemination into draining lymph nodes (dLN) in a CCR7-dependent manner[17]. Although i.d. infection can mimic the route of early bacterial dissemination, neither lethal outcomes nor skin eschars can be recapitulated in tested immunocompetent mice (B6, BABL/c, CD-1). It is therefore unclear as to how innate immune components or events successfully control bacterial dissemination and replication, and how such “gatekeepers” fail to protect the host.

Timely control of bacterial dissemination and replication is crucial to prevent systemic infection, hyperinflammation, and tissue damage in scrub typhus. However, it remains unclear as to how different strains of *Ot* bacteria react to or evade innate immunity in the early stages, and lead to different disease outcomes. We recently reported that i.v. inoculation with *Ot* Gilliam strain led to a self-limiting infection in B6 mice with restricted bacterial burdens in various organs as early as 4 days post-infection (p.i.)[18], implying Gilliam-induced protective immunity. In sharp contrast, *Ot* Karp strain can evade host innate immune defense, leading to uncontrolled bacterial dissemination and growth accompanied by severe pathology and high mortality rates[18]. Although CD8^+^ T cell-mediated responses contribute to *Ot* elimination[19, 20], dysregulated adaptive immune responses and excessive release of alarmin molecules (e.g., IL-33) are linked to hyperinflammation, vascular dysfunction, acute tissue damage, and host lethality[20–22]. Therefore, a better understanding of the initiation of innate immunity against *Ot* infection is urgently needed.

Prior microarray research of monocytes and MФs in human scrub typhus suggested that this infection triggers antiviral-like immune programs with elevated IFN-mediated responses[23, 24]. We also reported the extremely upregulated levels of IFN-γ, TNF-α, IFN-γ-regulated CXCR chemokines (CXCL9, CXCL10, CXCL11) in the lungs, spleen, liver, and brain of Karp-infected B6 and CD-1 mice[12, 25, 26]. Mouse brain RNAseq confirmed activated signaling pathways for IFN responses, defense response to bacteria, immunoglobulin-mediated immunity, the IL-6/JAK-STAT axis, and the TNF/NF-κB axis[27]. *Ot* Karp infection also induced substantial IFN-I signals and its associated genes (*Mx1*, *Oas1, Oas3*, etc.) in mouse brain and bone marrow-derived MФs[28]. Using *Rickettsia parkeri*, Burke et al. recently found that i.d. infection of mice lacking both IFN receptors (double *Ifnar1*^-/-^*Ifngr1*^-/-^) elicited eschar formation and lethal disease[29], suggesting the vital role of IFNs in Rickettsial infection[30, 31]. However, the functional role of IFN responses in scrub typhus is still elusive.

In the present study, we demonstrated that mice lacking either the IFN receptors (double *Ifngr1*^-/-^ *and Ifnar1*^-/-^) or STAT1 (the IFN-signaling downstream transcriptional factor) were highly susceptible to *Ot* infection, and exhibited skin lesions that were similar in appearance to human eschars. By using single knockout (*Ifnar1*^-/-^ or *Ifngr1*^-/-^) mice, we provided the first evidence for an essential role of IFN-γ, but not IFN-I, in host protection against *Ot* infection and skin eschar formation. By using IFN-γ reporter mice and neutralizing antibody, we provided evidence that NK cells were the major source of innate IFN-γ and contributed to early control of *Ot* infection. Notably, the relative contribution of IFN-γ signaling to eschar formation and animal survival is correlated with the *Ot* strain and infectious dose. Thus, this study provides new insights into the innate immunity and pathogenesis in a lethal infection mouse model with i.d. *Ot* inoculation.

## Materials and Methods

### Animals, infection, and treatment

B6 (#000664), double *Ifngr1*^-/-^*Ifnar1*^-/-^ (#029098), *Stat1*^-/-^ (#012606), *Ifngr1*^-/-^ (#003288) and IFN-γ reporter (#017581) mice were purchased from Jackson Laboratory.

*Ifnar1*^-/-^ (#032045-JAX) mice were purchased from Mutant Mouse Resource & Research Centers (MMRRC). Male mice were maintained under specific pathogen-free conditions and used at 8- 10 weeks of age, following protocols approved by the Institutional Animal Care and Use Committee (protocol # 2101001) at the University of Texas Medical Branch (UTMB) in Galveston, TX. All mouse infection studies were performed in the ABSL3 facility in the Galveston National Laboratory located at UTMB; all tissue processing and analysis procedures were performed in the BSL3 or BSL2 facilities. All procedures were approved by the Institutional Animal Care and Use Committee (IACUC) and the Institutional Biosafety Committee, in accordance with Guidelines for Biosafety in Microbiological and Biomedical Laboratories. UTMB operates to comply with the United States Department of Agriculture (USDA) Animal Welfare Act (Public Law 89–544), the Health Research Extension Act of 1985 (Public Law 99–158), the Public Health Service Policy on Humane Care and Use of Laboratory Animals, and the NAS Guide for the Care and Use of Laboratory Animals (ISBN-13). UTMB is a registered Research Facility under the Animal Welfare Act and has a current assurance on file with the Office of Laboratory Animal Welfare, in compliance with NIH Policy. For animal infection, mice were first anesthetized in a chamber connected to a VetFlo isoflurane vaporizer. Next, mice were carefully removed from the chamber and the infection site (right flank) was shaved using an electric trimmer. Mice were inoculated with *Ot* Karp or Gilliam strain (20 µL volume) in the dermis of the flank by using a 0.3 mL insulin syringe with 31G needle (Sol-Millennium, Chicago, IL), and were monitored for ∼5 minutes (min) until they were fully awake. Mice were monitored daily for weight loss, skin lesion, signs of disease, and survival. Some mice were intraperitoneally (i.p.) injected with Ultra-LEAF Purified anti-mouse NK-1.1 Antibody (clone#PK136; 200 µg/mouse, Biolegend, San Diego, CA) every other day, starting from 3 days prior to infection. Control mice were treated with Ultra-LEAF Purified Mouse IgG2a, κ Isotype Ctrl (200 µg/mouse). At indicated time-points, mice were euthanized by CO_2_ inhalation and tissues/blood were harvested for further analysis.

The disease severity score (ranged from 0-5) was based on an approved animal sickness protocol[18, 32]. The criteria included mobility/lethargy, hunching, fur ruffling, bilateral conjunctivitis, and weight loss: 0-normal behavior; 1-active, some weight loss (< 5%); 2-weight loss (6-10%), some ruffled fur (between shoulders); 3-weight loss (11-19%), pronounced ruffled fur, hunched posture, erythema, signs of reduced food/water intake; 4- weight loss (20-25%), decreased activity, bilateral conjunctivitis, incapable of reaching food/water; 5- non-responsive (or weight loss of greater than 25%) and animal needed to be humanely euthanized.

To evaluate skin lesion formation, the diameter of eschar lesion was measured by using a digital caliper rule. The mouse skin lesion was photographed, and the score was evaluated according to the publication associated with Rickettsial infection[29]. Briefly, 0- no visible lesion; 1-moderate redness; 2-extensive redness with lesion of 1-4 mm diam; 3-lesion of 4-10 mm diam; 4-lesion > 10 mm diam.

### Bacterial stock preparation

Bacteria were inoculated onto confluent monolayers in T150 cell culture flasks and gently rocked for two hours (h) at 37°C. After 2 h, Minimum Essential Medium (MEM) with 10% fetal bovine serum, 100 units/mL of penicillin and 100 µg/mL of streptomycin were added. Cells were harvested by scraping at day 7 p.i., re-suspended in MEM, and lysed using 0.5 mm glass beads and vortexing for 1 min. The cell suspension was collected and centrifuged at 300×g for 10 min to pellet cell debris and glass beads. The supernatant from one T150 flask was further inoculated onto new monolayers of five T150 flasks. This process was repeated for a total of six passages. Cells from five flasks were pooled in a 50 mL conical tube with 20 mL medium and 5 mL glass beads. The conical tubes were gently vortexed at 10 sec intervals for 1 min to release the intracellular bacteria and placed on ice. The tubes were then centrifuged at 700×g to pellet cell debris, and the supernatant was collected in Oakridge high speed centrifugation bottles, followed by centrifugation at 22,000×g for 45 min at 4°C to harvest bacteria. Sucrose-phosphate-glutamate buffer (0.218 M sucrose, 3.8 mM KH_2_PO_4_, 7.2 mM KH_2_PO_4_, 4.9 mM monosodium L-glutamic acid, pH 7.0) was used for preparing bacterial stocks, which were stored at -80°C[10, 12]. The Karp and Gilliam stocks were prepared in separated culture experiments to avoid potential contamination. The same lot of stocks were used for all experiments described in this study. The titers of the bacterial stocks were measured by focus forming assays, as previously described[13].

### qRT-PCR

Cells were lysed with RLT lysis buffer for 10 min, and RNA was extracted using RNeasy Mini kits (Qiagen, Germantown, MD), followed by the synthesis of cDNA using an iScript Reverse Transcription kit (Bio-Rad, Hercules, CA). Tissue samples were lysed using RLT buffer and 0.5 mm stainless steel beads and vortexing for 1 min. cDNA was amplified in a 10 μL reaction mixture containing 5 μL of iTaq SYBR Green Supermix (Bio-Rad, Hercules, CA) and 5 μM each of gene-specific forward and reverse primers. The PCR assays were denatured for 30 s at 95°C, followed by 40 cycles of 15 s at 95°C, and 60 s at 60°C, utilizing a CFX96 Touch real-time PCR detection system (Bio-Rad, Hercules, CA). Relative quantitation of mRNA expression was calculated using the 2^−ΔΔCt^ method. The primers are obtained from PrimerBank[33] and listed here. *Gapdh*: Forward 5’-AGGTCGGTGTGAACGGATTTG-3’, Reverse 5’- TGTAGACCATGTAGTTGAGGTCA-3’; *Ifnb*: Forward 5’- CAGCTCCAAGAAAGGACGAAC-3’, Reverse 5’-GGCAGTGTAACTCTTCTGCAT-3’; *Ifng*: Forward 5’-ATGAACGCTACACACTGCATC-3’, Reverse 5’- CCATCCTTTTGCCAGTTCCTC-3’.

### Quantitative PCR for Measuring Bacterial Burdens

To determine bacterial burdens, mouse tissues and cultured cells were collected and incubated with proteinase K. DNA was then extracted using a DNeasy Blood & Tissue Kit (Qiagen, Germantown, MD) according to the instructions, and used for qPCR assays, as previously described[17, 25, 27]. The 47-kDa gene was amplified using the primer pair OtsuF630 (5’- AACTGATTTTATTCAAACTAATGCTGCT-3’) and OtsuR747 (5’- TATGCCTGAGTAAGATACGTGAATGGAATT-3’) primers (IDT, Coralville, IA) and detected with the probe OtsuPr665 (5’-6FAM- TGGGTAGCTTTGGTGGACCGATGTTTAATCT-TAMRA) (Applied Biosystems, Foster City, CA) by SsoAdvanced Universal Probes Supermix (Bio-Rad, Hercules, CA). Bacterial burdens were normalized to total microgram (µg) of DNA per µL for the same samples. The copy number for the 47-kDa gene was determined by known concentrations of a control plasmid containing a single-copy insert of the gene. Gene copy numbers were determined via serial dilution (10-fold) of the *Ot* Karp 47-kDa plasmid.

### Bone marrow-derived DC generation and infection

Bone marrow-derived DCs were generated from B6 mice by cultivation with recombinant GM-CSF (20 ng/ml), as described previously[34]. Fresh GM-CSF-containing medium was added at days 3 and 6, and DCs were harvested at day 8 for seeding in 24-well plates. For in vitro infection, cells were infected with *Ot* Karp (MOI 10) for 2 h, followed by thoroughly washing with warm PBS. Fresh medium containing IFN-β or IFN-γ was added to the wells for continued culture in 37°C and 5% CO_2_. Cells were collected at 24, 48, 72, and 96 h p.i. for bacterial measurement. Mouse recombinant GM-CSF, IFN-β, and IFN-γ were purchased from Biolegend (San Diago, CA).

### Flow cytometry

For detection of infiltrated immune cells, lungs were perfused by injecting 10 mL PBS into the right ventricle of the heart. After perfusion, the whole left lobes were harvested and minced into small pieces, followed by the digestion with 0.05% collagenase type IV (Gibco/Thermo Fisher Scientific) in Dulbecco’s Modified Eagle’s Medium (DMEM, Sigma-Aldrich, St. Louis, MO) for 30 min at 37°C. Lung single-cell suspensions were made by passing lung homogenates through 70-μm cell strainers[10, 18]. Draining lymph nodes (dLN) and spleens were also collected and passed through cell strainers for preparing single cell suspensions[17]. Red blood cells were removed by using Red Cell Lysis Buffer (Sigma-Aldrich, St. Louis, MO) for 5 min at room temperature. For surface marker analysis, leukocytes were stained with the Fixable Viability Dye (eFluor 506, Thermo Fisher Scientific, Waltham, MA) for live/dead cell discrimination, blocked with FcγR blocker, and incubated with fluorochrome-labeled antibodies for 30 min. The fluorochrome-labeled antibodies were purchased from Thermo Fisher Scientific and Biolegend as below: Alexa Fluor 700 anti-CD11b (M1/70), APC anti-Ly6G (1A8), PE/Dazzle-594 anti-Ly6C (HK1.4), PE anti-CD64 (X54-5/7.1), Percp-Cy5.5 anti-CD11c (N418), FITC anti-MHCII (M5/114.15.2), PE-Cy7 anti-CD3ε (145-2C11), Percp-cy5.5 anti-CD4 (GK1.5), APC-Cy7-anti-CD8a (53–6.7), BV711 anti-CD44 (IM7), APC anti-CD62L (MEL-14), PE CF594-anti-NK1.1 (PK136) and FITC anti-CD69 (H1.2F3). BV421 anti-F4/80 (T45-2342) antibody was purchased from BD Bioscience (San Diago, CA). Cell samples were fixed in 2% paraformaldehyde overnight at 4°C, acquired by a BD LSR Fortessa and analyzed via FlowJo software version 10 (BD, Franklin Lakes, NJ). The gating strategy of immune cell subsets are based on recent publications[10, 18, 35].

### Histology

Tissues were fixed in 10% neutral buffered formalin and embedded in paraffin at the UTMB Research Histology Service Core. Tissue sections (5-μm thickness) were stained with hematoxylin and eosin and mounted on slides. Sections were imaged under an Olympus BX53 microscope, and at least five random fields for each section were captured.

### Bio-Plex assay

Whole blood was collected from anesthetized mice at various timepoints, and serum was isolated by using serum separator tubes (BD Bioscience, San Diago, CA), followed by the bacterial inactivation, as described in our previous studies[12, 13, 32]. The customer-designed Bio-Plex kits were used to measure the levels of serum cytokines and chemokines (IL-2, IL-4, IL-5, IL-6, IL-10, IL-12p40, IL-13, IL-17, G-CSF, IFN-γ, CCL1, CCL2, CCL3, CCL4, CCL5, CCL11, CXCL1, CXCL10, CXCL12, and CXCL16). The Bio-Rad Bio-Plex Plate Washer and Bio-Plex 200 machines were used for sample processing and analysis in UTMB Flow Cytometry and Cell Sorting Core Lab.

### Clinical pathology

Animal blood chemistry analysis was performed by using the VetScan Chemistry Analyzer (Zoetis, Parsippany-Troy Hills, NJ), according to the manufacturer’s instruction. Briefly, mouse serum (100 μL) was uploaded into the VetScan Comprehensive Diagnostic Profile reagent rotor, which was used for quantitative determinations of alanine aminotransferase (ALT), albumin (ALB), alkaline phosphatase (ALP), amylase (AMY) total calcium (CA^++^), creatinine (CRE), globulin (GLOB), glucose (GLU), phosphorus (PHOS), potassium (K^+^), sodium (NA^+^), total bilirubin (TBIL), total protein (TP), and urea nitrogen (BUN). This analysis was performed in UTMB ABSL3 animal facility.

### Statistical analysis

Data are presented as mean ± standard deviation (SD). qRT-PCR and some bacterial burden data were analyzed with one-way ANOVA. For other data including body weight changes, disease scores, skin lesion scores, flow cytometry, serum Bio-Plex and chemistry parameters, two-way ANOVA was used for statistical analysis. After a significant F-test for the ANOVA model, either Tukey’s multiple comparisons test or Šídák’s multiple comparisons test was used for comparison between groups. Survival curves were analyzed by using the Log-rank (Mantel-Cox) test. All data were analyzed by using GraphPad Prism software 10. Statistically significant values are denoted as * *p* < 0.05, ** *p* < 0.01, *** *p* < 0.001, and **** *p* < 0.0001, respectively.

## Results

### Mice lacking IFN signals are highly susceptible to intradermal *Ot* Karp infection

IFN and downstream signaling are considered the critical components of innate and adaptive immune defense against viral and bacterial infections. Our previous studies have demonstrated the robust type I immunity in various vesicle organs in the i.v. mouse models of *Ot* infection[12, 22, 25–27, 36]. Here, we sought to utilize the i.d. murine infection model to better recapitulate the natural route of *Ot* infection and investigate the role of IFN signals. As shown in **Fig. 1A and B**, both *Ifng* and *Ifnb* transcripts were significantly increased in the skin and dLN at day 1 p.i.. IFN-γ levels were markedly upregulated by 100-fold in the dLN on day 3 p.i., compared to the mock group (**Fig. 1B**). These results indicate that *Ot* induces early IFN responses at the infection site. To uncover the role of IFN signaling in *Ot* infection in the initial bacterial, we infected WT B6, double *Ifnar1*^-/-^*Ifngr1*^-/-^, and *Stat1*^-/-^ mice with *Ot* Karp. As we previously reported[16, 17], i.d. infection in B6 WT mice resulted in a non-lethal infection characterized by limited body weight changes (∼5-10%) and mild disease symptoms (score < 2) (**Fig. 1C-E**). However, mice lacking IFN receptors or the downstream transcriptional factor STAT1 were highly susceptible to *Ot* infection, as evidenced by dramatic body weight loss (∼15-20%) and severe disease symptoms (score > 3) on days 10 or 11 p.i. (**Fig. 1C-E**). Both types of gene-deficient mice exhibited significantly higher bacterial burdens in tested organs (spleen, lung, liver, kidney, and brain), and succumbed before day 13 p.i. (**Fig. 1E-F**). Histological assessment showed visible lung edema and obvious liver necrosis on day 10 p.i. (**Fig. 1G**), indicating that severe pathology results from *Ot* infection in the absence of IFN signaling. During our daily monitoring, no dermal lesions appeared at the site of infection in WT mice. In contrast, both double *Ifnar1*^-/-^ *Ifngr1*^-/-^ and *Stat1*^-/-^ mice displayed evident skin lesions marked by necrosis of the overlying epidermis and hardening features, surrounded by an indurated red halo (**Fig. 1H, red arrows**), which resembled skin eschars in scrub typhus patients[37–39]. Therefore, the i.d. infection in mice lacking IFN signals reproduces the characteristic manifestations observed in human *Ot* infection, suggesting a critical role of IFNs in disease control and skin eschar formation in scrub typhus.

**Fig. 1.**
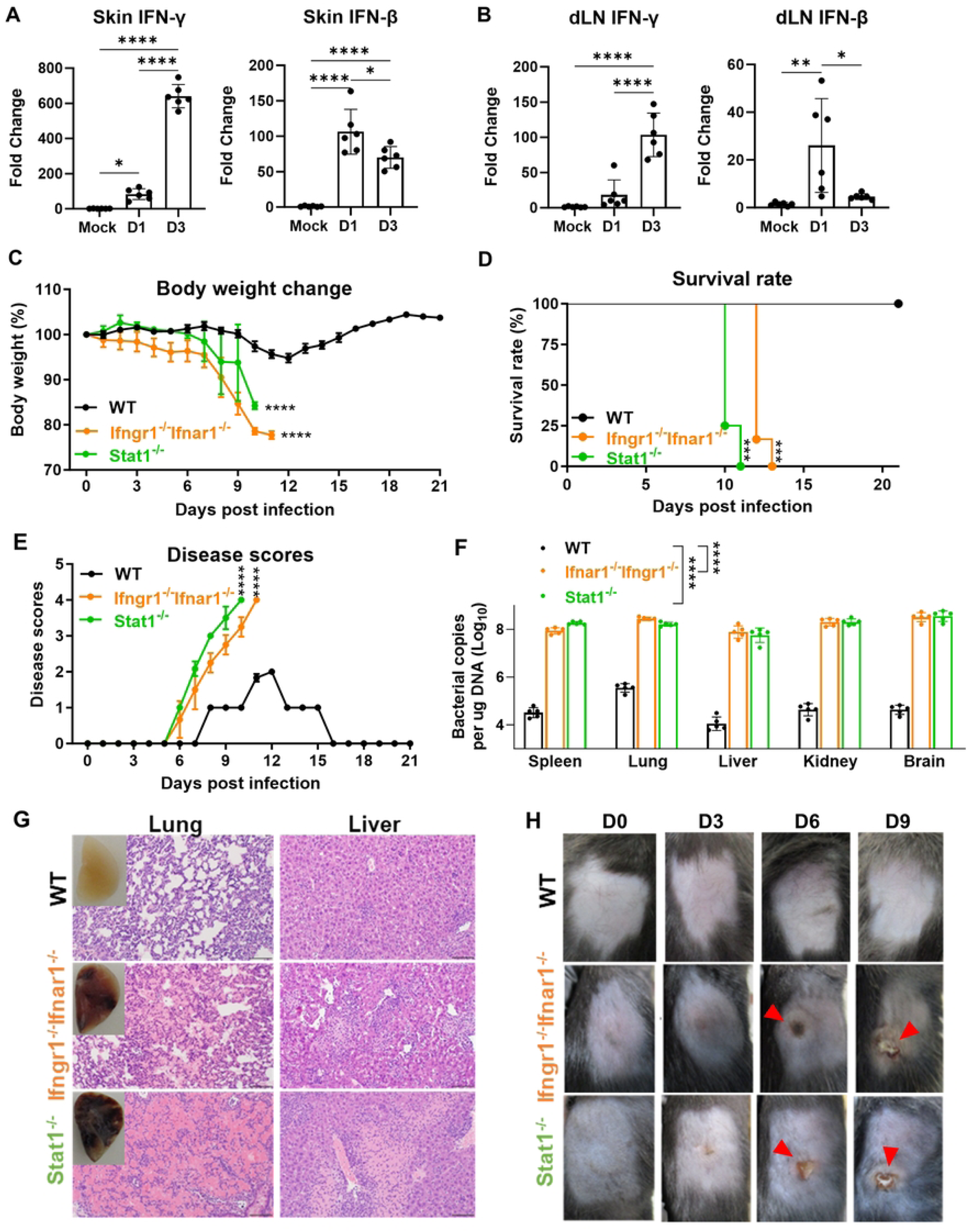
Mice lacking both interferon signals (double *Ifnar1*^-/-^*Ifngr1*^-/-^) are highly susceptible to intradermal *Ot* Karp infection. B6 mice (n = 6/group) were intradermally (i.d.) infected with *Ot* Karp strain (3×10^3^ FFU) on the flank. (A) Skin and (B) draining lymph nodes (dLN) were harvested and lysed by RLT buffer using stainless steel beads for vortexing. RNA was extracted and cDNA was synthesized for qRT-PCR of IFN-γ and IFN-β. (C) WT B6, double *Ifnar1*^-/-^*Ifngr1*^-/-^ and *Stat1*^-/-^ mice (n = 5/group) were i.d. infected with *Ot* Karp strain (3×10^3^ FFU) in the flank. Body weight changes, (D) survival rates and (E) disease scores were monitored daily. (F) Bacterial loads were measured in the spleen, lung, liver, kidney, and brain. The results of statistical analysis represent the comparisons for all five organs. (G) The representative histological images of the lung and liver on day 10 p.i. are shown. The photos of perfused lungs are also included. Scale bar = 200 µm. (H) The photos of skin inoculation sites at days 0, 3, 6 and 9 p.i. are presented. The skin eschar-like lesions are indicated by red arrows. The values are shown as mean ± SD from single experiments and are representative of two independent experiments. A one-way ANOVA with a Tukey’s multiple comparisons test was used for statistical analysis of qRT-PCR and bacterial load data. Body weight changes and disease scores were analyzed by two-way ANOVA and Šídák’s multiple comparisons test. Survival curves were analyzed by using Log-rank (Mantel-Cox) test. *, *p* < 0.05; **, *p* < 0.01; ***, *p* < 0.001; ****, *p* < 0.0001.

### IFN-γ, but not IFN-I, plays a key role for host protection against *Ot* Karp infection

While both IFN-γ and IFN-I can activate the JAK/STAT pathway and exhibit similar abilities to ‘interfere’ with viral infections, they regulate distinct effector cells based on the binding of specific receptors[40]. To examine the individual role of each IFN signal, we infected WT, *Ifnar1*^-/-^ and *Ifngr1*^-/-^ mice with *Ot* Karp. As shown in **Fig. 2A-C**, *Ifngr1*^-/-^ mice were highly susceptible to infection, as demonstrated by significant body weight loss (> 20%) and higher disease symptoms (score = 4) on day 12 p.i.. All *Ifngr1*^-/-^ mice succumbed around day 14 p.i. and demonstrated similar disease severity to that observed in double *Ifnar1*^-/-^*Ifngr1*^-/-^ mice (Fig. 1). In contrast, all WT and *Ifnar1*^-/-^ mice survived and were active with limited body weight loss (< 10%) and disease symptoms (score = 1-2). More importantly, *Ifngr1*^-/-^ mice, but not WT or *Ifnar1*^-/-^ mice, developed an eschar lesion at the infection site starting from day 6 p.i. (**Fig. 2D**). In line with that, histological analysis revealed extensive inflammatory infiltration in the lungs and liver of *Ifngr1*^-/-^ mice, along with significant hepatocyte necrosis on day 10 p.i. (**Fig. 2E**). We also evaluated animal health status by monitoring mouse serum chemistry indicators. As described in **Fig. 2F**, *Ifngr1*^-/-^ mice exhibited higher levels of total protein (TP) and globulin (GLOB) at days 6 and 10 p.i., indicating a more pronounced inflammatory response to infection. The lack of IFN-γ signaling also caused liver dysfunction and malnutrition, as evidenced by decreased serum albumin (ALB) and glucose (GLU) levels. Decreased amylase (AMY), alanine aminotransferase (ALT), and alkaline phosphatase (ALP) were also found in *Ifngr1*^-/-^ mice on days 6 and 10 p.i., as compared to WT mice, suggesting that the regulation of these enzymes might be attributed to IFN-γ signals in the later. Furthermore, the elevated levels of serum sodium (Na^+^), potassium (K^+^), and calcium (Ca^++^) observed in *Ifngr1*^-/-^ mice only on day 10 p.i. suggested the presence of severe dehydration at the peak of disease. No significant difference was observed for creatinine (CRE), phosphorus (PHOS), total bilirubin (TBIL), and urea nitrogen (BUN) between WT and *Ifngr1*^-/-^ mice at different time points (**Fig. S1**). Thus, we provided the strong evidence that IFN-γ, but not IFN-I, was vital for host immune protection and skin eschar formation following *Ot* infection.

**Fig. 2.**
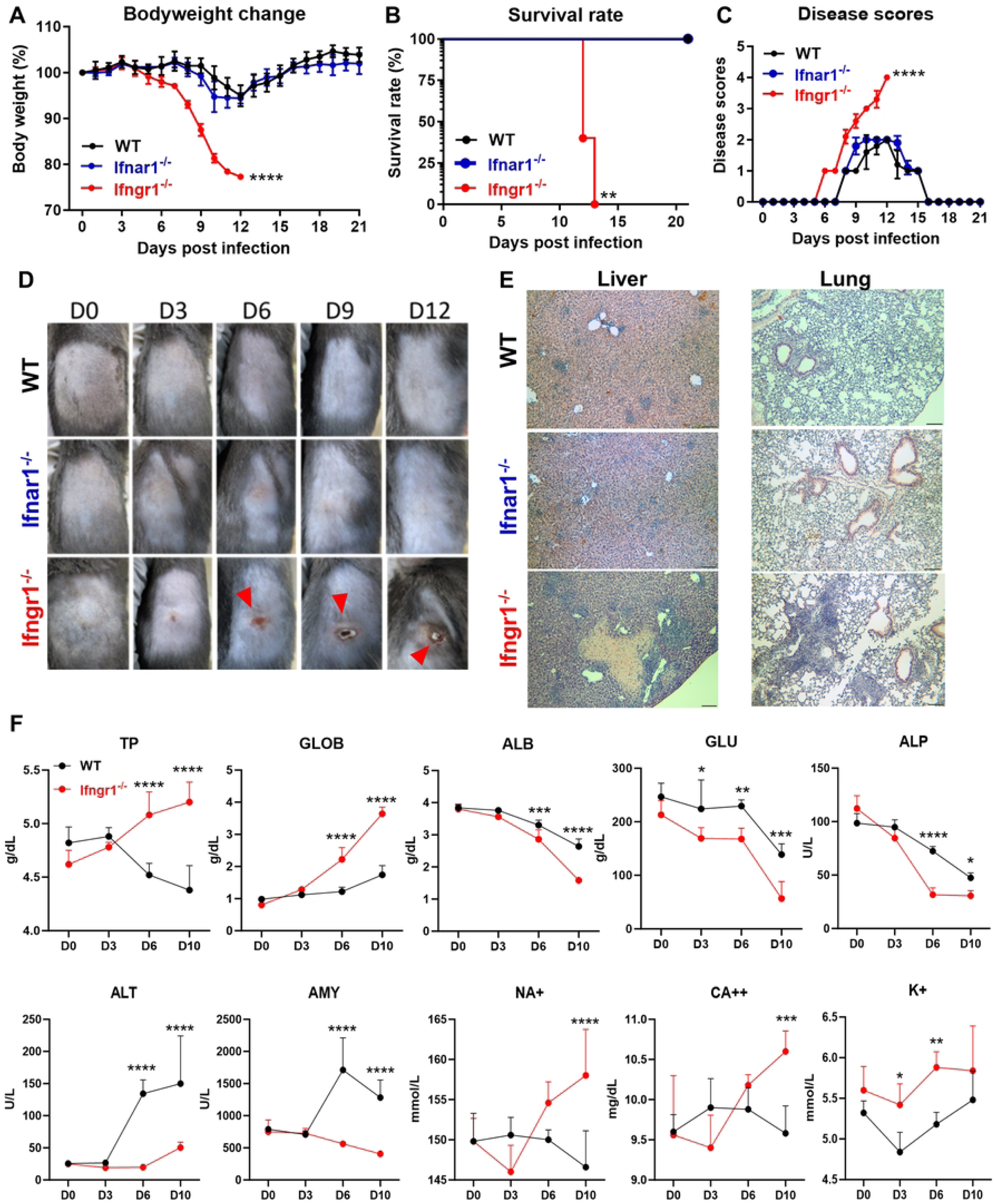
IFN-γ, but not IFN-I is required for host protection against *Ot* Karp infection. WT, Single *Ifnar1*^-/-^ *and Ifngr1*^-/-^ (n = 5/group) were i.d. infected with *Ot* Karp strain (3×10^3^ FFU) on the flank. (A) Body weight changes, (B) survival rates and (C) disease scores were monitored daily. (D) The photos of skin inoculation sites at days 0, 3, 6, 9 and 12 p.i. are presented. The skin eschar-like lesions are indicated by red arrows. (E) The representative histological images of the lung, liver and brain are shown. Scale bar = 200 µm. (F) The mouse serum chemistry profile was generated by using VetScan Comprehensive Diagnostic Profile reagent rotor. The parameters include alanine aminotransferase (ALT), albumin (ALB), alkaline phosphatase (ALP), amylase (AMY) total calcium (CA^++^), globulin (GLOB), glucose (GLU), potassium (K^+^), sodium (NA^+^), total protein (TP). The values are shown as mean ± SD from single experiments and are representative of two independent experiments. Body weight changes, disease scores, and serum chemistry profile were analyzed by two-way ANOVA and Šídák’s multiple comparisons test. Survival curves were analyzed by using Log-rank (Mantel-Cox) test. *, *p* < 0.05; **, *p* < 0.01; ***, *p* < 0.001; ****, *p* < 0.0001.

### Immune dysregulation in the absence of the IFN-γ signal during *Ot* infection

We next dissected immune responses in the absence of the IFN-γ signal in *Ot* infection. Since the lung is the primary organ targeted by *Ot*, we extracted immune cells infiltrating the lungs from both WT and *Ifngr1*^-/-^ mice for flow cytometric analysis. We examined both myeloid and lymphocyte subpopulations (**Fig. 3A**) and found a significantly increased number of neutrophils in the *Ifngr1*^-/-^ mice on days 6 and 10 p.i. (**Fig. 3B**). Inflammatory monocytes, which were characterized as the Ly6C^hi^ population also showed a 17-fold increase on day 6 p.i. and a 57-fold increase on day 10 p.i. in *Ifngr1*^-/-^ mice compared to those in WT mice (**Fig. 3B**). Together, these findings suggest that both neutrophils and monocytes may contribute to exacerbated lung pathology in *Ot* infection. In contrast, the number of total MФ, alveolar MФ (AMФ), and interstitial MФ (IMФ) were greatly reduced in *Ifngr1*^-/-^ mice on both days 6 and 10 p.i., indicating the necessity of IFN-γ for maintaining pulmonary MФ responses. In addition, the absence of IFN-γ did not cause a deficiency of adaptive T cell response; instead, it led to an increased infiltration of activated T cells in the lung on day 10 p.i. (**Fig. 3C**). Similarly, the number of activated CD4 and CD8 T cells were comparable in the spleens (**Fig. S2**). In contrast, the number of activated NK cells were dramatically decreased in both spleen and lungs on days 6 and 10 p.i. (**Fig. 3C and S2**), indicating the key role of IFN-γ in NK cell activation. Together, our flow cytometry data demonstrate a dysregulated immune response to *Ot* infection in the absence of the IFN-γ signal.

**Fig. 3.**
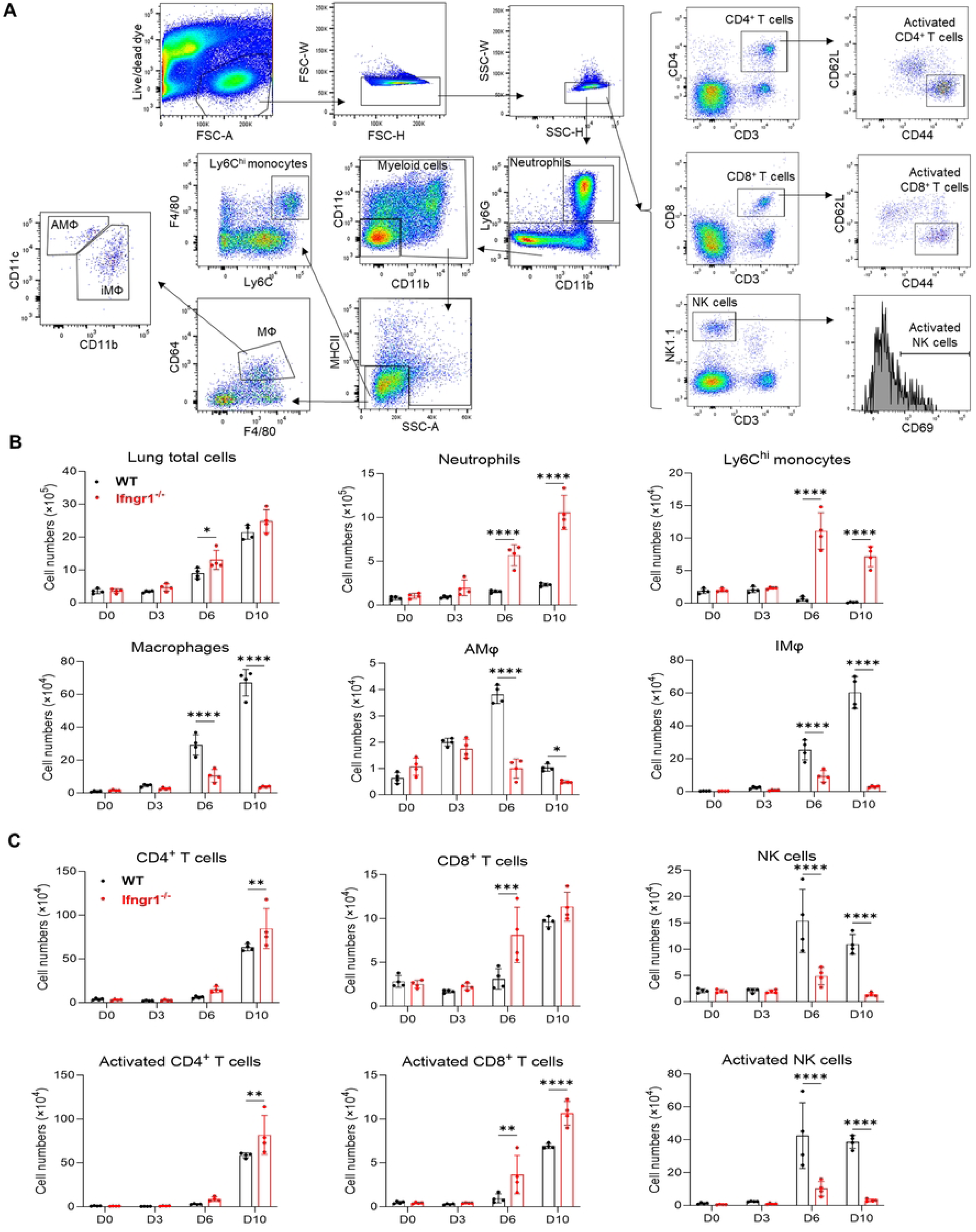
Dysregulated lung immune responses in the absence of the IFN-γ signal during *Ot* infection. WT and *Ifngr1*^-/-^ mice (n = 4/group) were i.d. infected with *Ot* Karp (3×10^3^ FFU). Mouse lungs were perfused with cold PBS and collected for preparation of single cell suspensions at days 3, 6 and 10 p.i.. Mock-infected mice were used at day 0. Cell samples were acquired by flow cytometry and data were analyzed by using Flowjo. (A) Gating strategy of lung immune cells is shown. Dead cells and doublets were first excluded by live/dead fixable dye and FSC-H vs FSC-W/SSC-H vs SSC-W, respectively. The live and single cells were gated for neutrophils (CD11b^+^Ly6G^+^), monocytes (Ly6G^-^MHCII^-^SSC^lo^F4/80^+^Ly6C^hi^), alveolar macrophages (AMФ, Ly6G^-^MHCII^-^SSC^lo^ CD64^+^F4/80^+^CD11c^+^CD11b^-^), interstitial macrophages (IM, Ly6G^-^MHCII^-^SSC^lo^ CD64^+^F4/80^+^CD11c^-^CD11b^+^), CD4 T cells (CD3^+^CD4^+^), CD8 T cells (CD3^+^CD8^+^), NK cells (CD3^-^NK1.1^+^), activated CD4 T cells (CD3^+^CD4^+^CD44^+^CD62L^-^), activated CD8 T cells (CD3^+^CD8^+^CD44^+^CD62L^-^), and activated NK cells (CD3^-^NK1.1^+^CD69^+^). (B) Cell numbers of total lung tissues, neutrophils, monocytes, and macrophage subsets are shown. (C) Cell numbers of CD4 T cells, CD8 T cells, NK cells, and their respective activated subsets in the lung are depicted. The values are shown as mean ± SD from single experiments and are representative of two independent experiments. Two-way ANOVA was used for statistical analysis. Šídák’s multiple comparisons test was used for multiple comparisons between WT B6 and *Ifngr1*^-/-^ mice at each time. *, *p* < 0.05; **, *p* < 0.01; ***, *p* < 0.001; ****, *p* < 0.0001.

### Dynamic serum cytokine/chemokine profile of WT and *Ifngr1*^-/-^ mice by *Ot* infection

To systematically evaluate immune status in WT and *Ifngr1*^-/-^ mice during *Ot* infection, we assessed dynamic serum cytokine/chemokine levels by Bio-Plex assay. As shown in **Fig. 4A**, the production of IL-2 and IL-12p40 (the essential cytokines for T cell activation and proliferation) were lower in *Ifngr1*^-/-^ mice. Likewise, TNF-α (the critical cytokine known for host protection against *Ot* infection[32]) was also reduced in *Ifngr1*^-/-^ mice on days 6 and 10 p.i.. IL-10 (an anti- inflammatory cytokine) was also decreased in the deficient mice on day 6 p.i. as compared to WT mice. These results indicate that Ifngr1 deficiency impairs the balance between pro-and anti-inflammatory cytokine response. In addition, Th2 cytokines including IL-4 and IL-5 were also decreased in *Ifngr1*^-/-^ mice following infection (**Fig. 4A**), suggesting the downregulation of both type 1 and 2 immune responses in the absence of IFN-γ signaling. Moreover, *Ifngr1*^-/-^ mice exhibited lower levels of chemokines, including CCL3, CCL5, CCL11, CXCL1, CXCL10, and CXCL16, which are important for the migration of activated T cells, MФ and neutrophils (**Fig. 4A**). In contrast to the downregulated cytokines and chemokines, IL-6 and G-CSF were significantly increased in *Ifngr1*^-/-^ mice on days 6 and 10 p.i. (**Fig. 4B**). Both IL-6 and G-CSF, mainly produced by myeloid cells and endothelial cells are potent inflammatory cytokines and may contribute to pathogenesis in *Ot* infection[18, 41]. CCL1 acts as a potent attractant for Th2 cells and a subset of T-regulatory (Treg) cells[42, 43]. CXCL12 is also involved in the maintenance of the Th2 cell function and was previously found to be downregulated in our scrub typhus mouse model[12, 44]. As shown in **Fig. 4B**, increased levels of CCL1 and CXCL12 were observed in *Ifngr1*^-/-^ mice on days 6 p.i., indicating that the lack of IFN-γ signal resulted in elevated Th2 chemokines in *Ot* infection. *Ifngr1*^-/-^ mice also exhibited a high level of compensatory IFN-γ production following *Ot* infection (**Fig. 4B**). No significant difference was found for IL-17, CCL2, and CCL2 between WT and *Ifngr1*^-/-^ mice during infection (**Fig. 4C**).

**Fig. 4.**
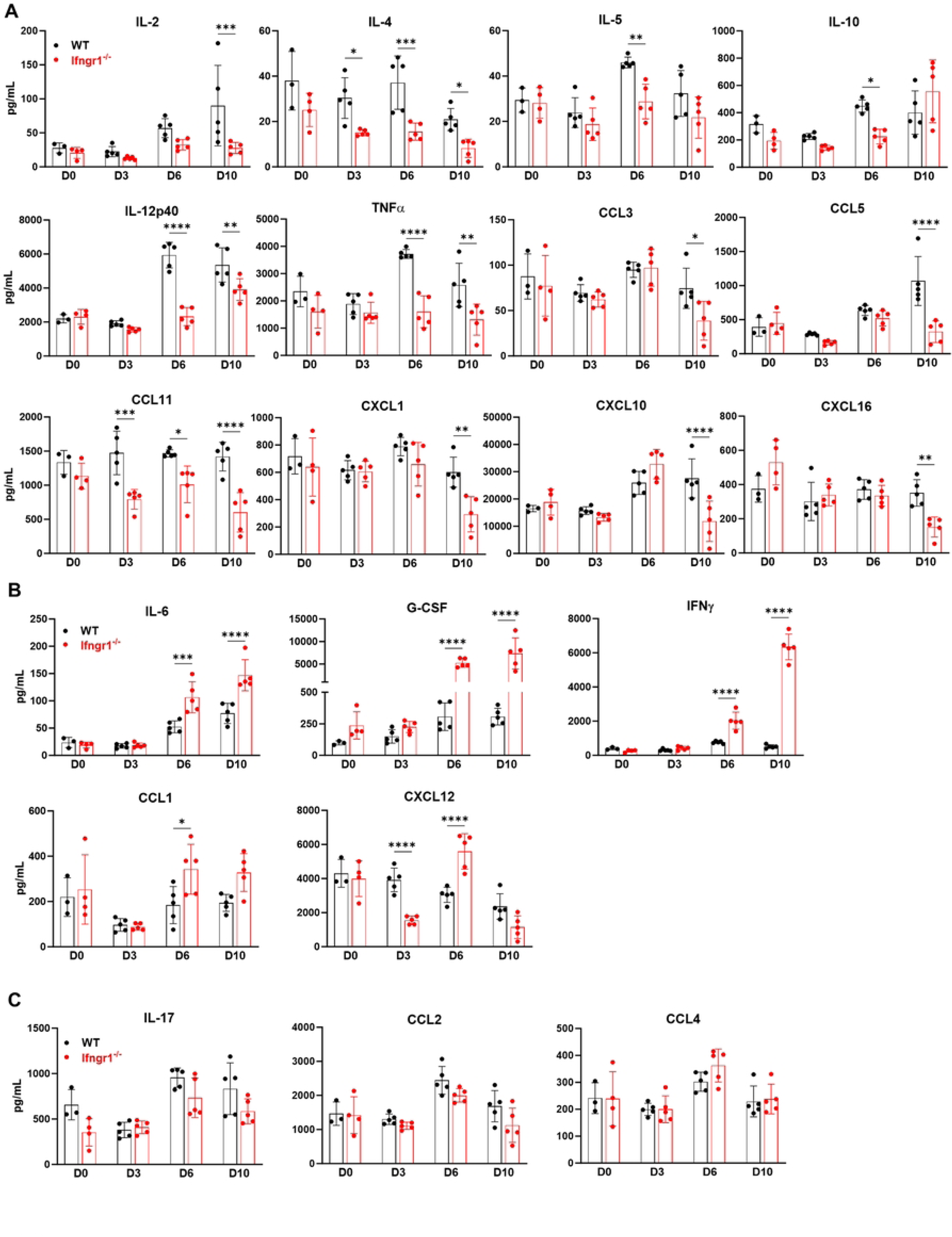
Dynamic systemic cytokine/chemokine responses in WT B6 and *Ifngr1*^-/-^ mice following *Ot* infection. WT and *Ifngr1*^-/-^ mice (n = 5/group) were i.d. infected with *Ot* Karp (3×10^3^ FFU). Mouse blood was collected at different time points, and serum was separated by using BD serum separator tubes. Serum cytokines/chemokines were measured by Bio-Rad Bio- Plex assay. As compared to WT mice, (A) *Ifngr1*-deficiency resulted in decreased production of IL-2, IL-4, IL-5, IL-10, IL-12p40, TNF-α, CCL3, CCL5, CCL11, CXCL1, CXCL10, and CXCL16. (B) *Ifngr1*-deficiency resulted in decreased production of IL-6, G-CSF, IFN-γ, CCL1, and CXCL12. (C) No differences were found for IL-17, CCL2, and CCL4 between WT and *Ifngr1*^-/-^ mice. The values are shown as mean ± SD from single experiments and are representative of two independent experiments. Two-way ANOVA was used for statistical analysis. Šídák’s multiple comparisons test was used for multiple comparisons between WT B6 and *Ifngr1*^-/-^ mice at each time. *, *p* < 0.05; **, *p* < 0.01; ***, *p* < 0.001; ****, *p* < 0.0001.

### IFN-γ signal is critical for bacterial control during *Ot* infection

To determine how IFN signaling contributes to the control of *Ot* growth, we measured the bacterial burdens in various organs (**Fig. 5A**). We observed significantly higher bacterial burdens in *Ifngr1*^-/-^ mice across all detected organs on day 6 p.i. compared to WT mice. By day 10, the bacterial burdens in organs had increased to approximately 100-fold higher in *Ifngr1*^-/-^ mice. No differences in bacterial burdens were observed in the lung and liver of *Ifnar1*^-/-^ mice on day 10 p.i. as compared to WT mice (**Fig. S3**). Unlike other organs, the dLN of *Ifngr1*^-/-^ mice exhibited 30-fold higher bacterial burdens on day 3 p.i. as compared to those in WT mice (**Fig. 5A**). This result suggested that innate IFN-γ play a role in controlling bacterial dissemination or replication in dLN. Since we recently reported that DCs are the major target for *Ot* dissemination from the skin into the dLN[17], we sought to determine whether IFN-γ or IFN-I can inhibit *Ot* growth in DCs. To test this, we infected bone marrow-derived DCs with *Ot* (MOI 10) in the presence of recombinant IFNs, and measured the bacterial burden at various time points. As presented in **Fig. 5B**, IFN-β failed to inhibit bacterial growth at 24, 48 and 72 h regardless of the doses, whereas high-dose of IFN-γ (10,000 U/mL) inhibited bacterial growth at 24 h. Consistently, IFN-γ significantly controlled bacterial growth on both 48 and 72 h at varying doses. IFN-β contributed to bacterial control only at 96 h, although its inhibitory effect was not as potent as that of IFN-γ at the same applied doses. Therefore, our in vivo and in vitro data demonstrated that IFN-γ, plays a key role in bacterial control.

**Fig. 5.**
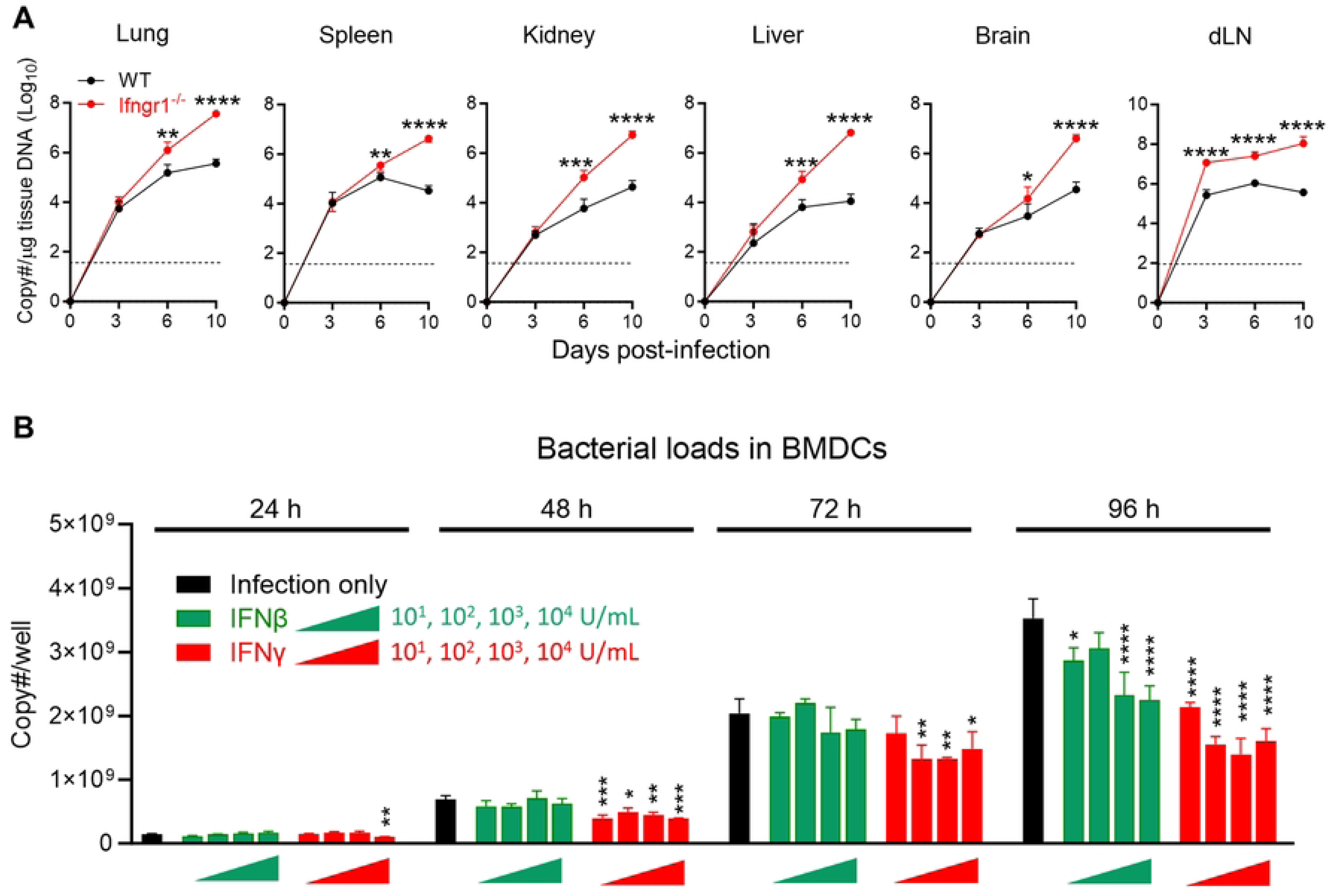
The deficiency of *Ifngr1* led to bacterial outgrowth in various organs after *Ot* infection. (A) WT and *Ifngr1*^-/-^ mice (n = 5/group) were i.d. infected with *Ot* Karp (3×10^3^ FFU). Mice were euthanized at days 3, 6, and 10 p.i., and mock-infected mice were used as a control (day 0). Mouse tissues (dLN, spleen, lung, liver, kidney, and brain) were harvested for bacterial burden analyses. (B) Bone marrow-derived DCs were infected with *Ot* Karp (MOI 10) in the presence of IFN-β or IFN-γ at indicted concentrations (10, 100, 1000, 10000 U/mL). This experiment was performed by triplicates. Bacterial burdens in the wells were measured on days 1, 2, 3 and 4 p.i. by qRT-PCR. The values are shown as mean ± SD from single experiments and are representative of two independent experiments. Two-way ANOVA was used for statistical analysis and Šídák’s multiple comparisons test was used for multiple comparisons between WT B6 and *Ifngr1*^-/-^ mice at each time. A one-way ANOVA statistical analysis with a Tukey’s multiple comparisons test was used for multiple comparisons of each infected group to the mock group at each time point. *, *p* < 0.05; **, *p* < 0.01; ***, *p* < 0.001; ****, *p* < 0.0001.

### NK cells are the main source of IFN-γ in the skin and dLN at early *Ot* infection

We demonstrated that innate IFN-γ contributed to bacterial control and may determine the subsequent immune activation against *Ot* infection in Fig. 4; however, a better understanding of this innate immunity would be advantageous for identifying potential therapeutic targets. We thus examined the cellular source of IFN-γ at early stages. To achieve this, we infected IFN-γ reporter mice and analyzed the IFN-γ^+^ cells in the skin and dLN, respectively. We found that both T cells (CD4^+^ and CD8^+^) and myeloid cells (CD11b^+^) in the dLN produced IFN-γ on days 1 and 3 p.i., with the positive percentages remaining below 6% (**Fig. 6A**). However, the percentages of IFN-γ^+^ NK cells were much higher than those of T cells and myeloid cells, reaching approximately 50% in the dLN on day 3 p.i.. *Ot* infection also significantly increased percentages of NK cells and IFN-γ-expressing NK cells in the skin at the early stage of infection (**Fig. 6B**). Thus, we demonstrated that NK cells were the major source of IFN-γ in the skin/dLN axis during early infection.

**Fig. 6.**
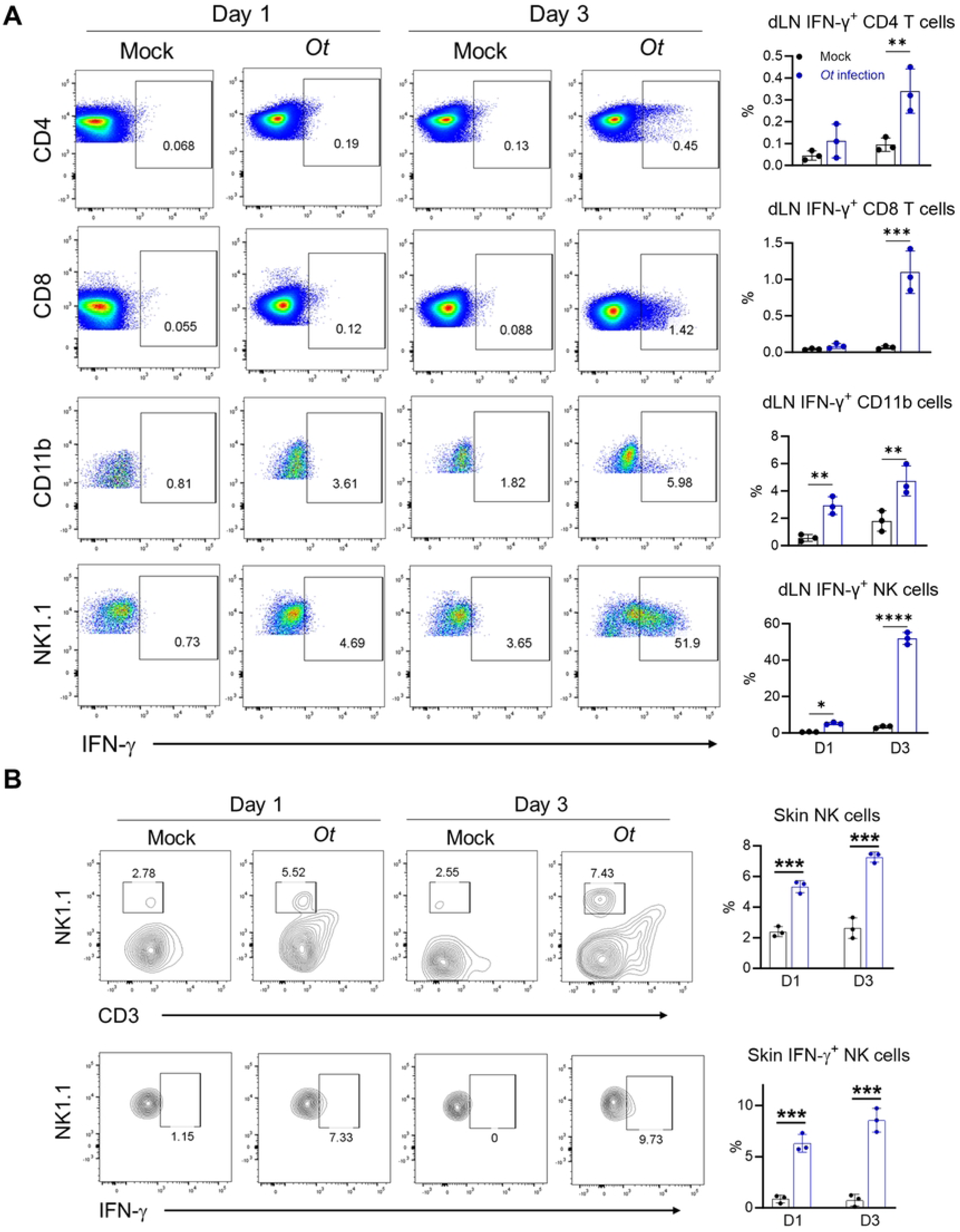
NK cells are the main source of IFN-γ at the early stage of *Ot* infection. IFN-γ reporter mice (n = 3/group) were i.d. infected with *Ot* Karp (3×10^3^ FFU). Mock-infected mice were used as controls. Mice were euthanized at days 1 and 3 p.i.. Mouse skin and dLN were harvested for flow cytometric analysis. (A) IFN-γ levels in CD4, CD8, myeloid, and NK cells of dLN were analyzed. (B) Percentage of NK cells and IFN-γ-expressing NK cells were analyzed in the skin. The values are shown as mean ± SD from single experiments and are representative of two independent experiments. Two-way ANOVA was used for statistical analysis. Šídák’s multiple comparisons test was used for multiple comparisons between mock and *Ot*-infected mice at each time. *, *p* < 0.05; **, *p* < 0.01; ***, *p* < 0.001; ****, *p* < 0.0001.

### NK cells contribute to early bacterial control

To confirm the functional role of NK cells, we depleted NK cells in *Ot*-infected mice by i.p. injection with an anti-NK1.1 antibody starting from -3 days prior to infection (**Fig. 7A**). While this depletion regimen greatly reduced the total number of NK cells in the dLN, lung, and spleen (**Fig. S4**), it did not cause a lethal infection, as neutralizing antibody-treated mice only showed slightly body weight loss on days 13 and 14 p.i. compared to IgG-treated mice (**Fig. 7B**). Of note, NK cell depletion resulted in increased bacterial burdens in the dLN on days 3 and 6 p.i. (**Fig. 7C**), but not in the lung and liver across all four time points (days 3, 6, 10 and 14 p.i.). To further evaluate the inflammatory responses in the absence of NK cells, we measured serum cytokines and chemokines levels using Bio-Plex assays. As shown in **Fig. 7D**, NK cell depletion resulted in decreased IL-5, IL-10, and G-CSF only on day 6 p.i.. Interestingly, TNF-α and IL-17, which are typical pro-inflammatory cytokines in human scrub typhus[12, 45, 46], were increased on days 10 and 14 p.i., respectively (**Fig. 7E**). We did not find any differences of other tested 14 cytokines and chemokines in our assay (**Fig. S5**). Therefore, our data demonstrated the role of NK cells in controlling bacteria in early *Ot* infection.

**Fig. 7.**
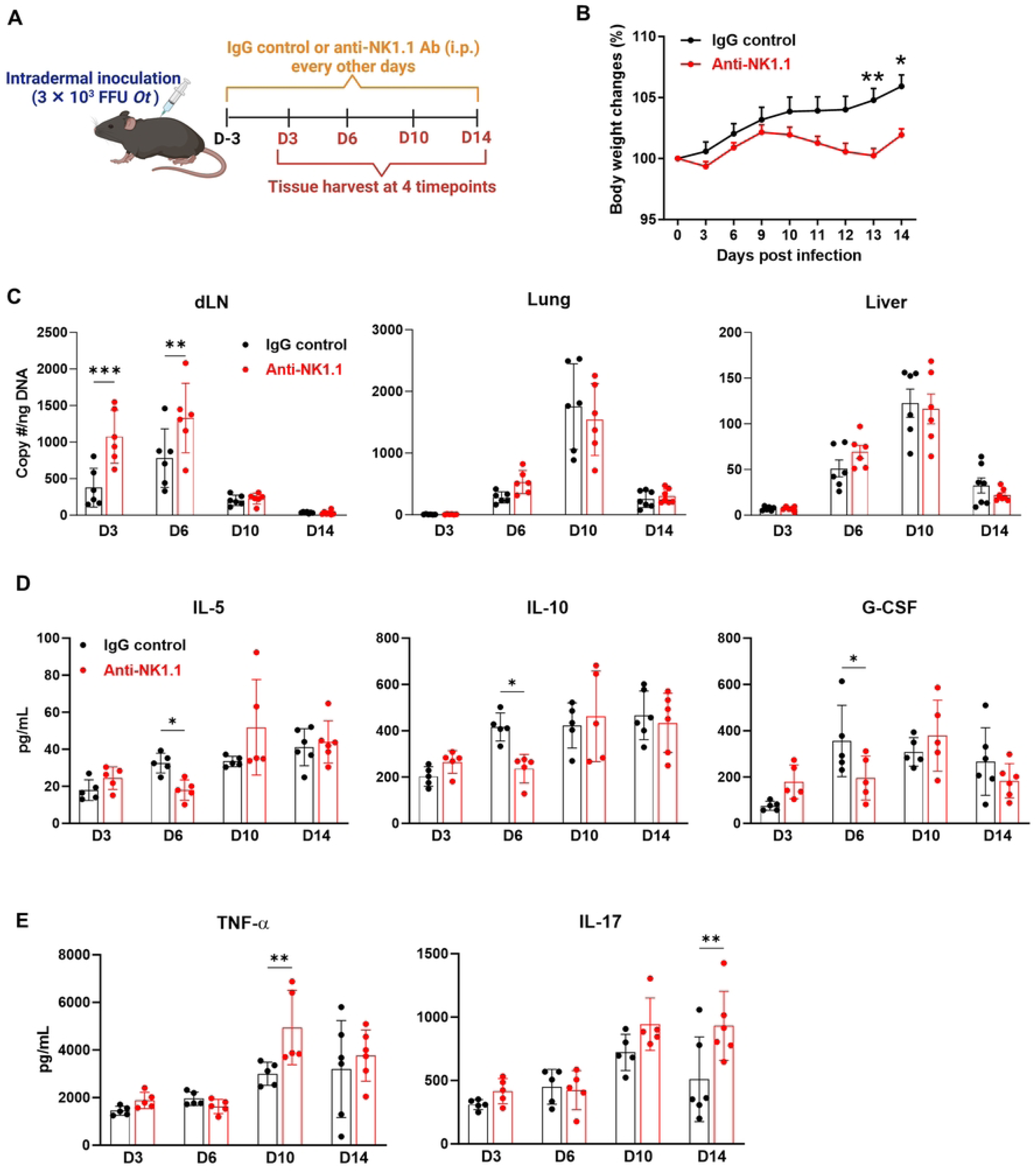
NK cells contribute to early bacterial control in dLN. (A) B6 mice (n = 5-6/group) were i.d. infected with *Ot* Karp (3×10^3^ FFU) and were i.p. treated with either IgG control or anti- NK1.1 neutralizing antibody (200 µg/mouse/treatment) every other day starting from 3 day prior to infection. Mice were euthanized at days 3, 6, 10, and 14 p.i. for tissue/blood harvest. (B) Body weight changes were monitored daily. (C) Bacterial burdens were measured in the dLN, lung, and liver at various time points. (D-E) Mouse serum was assessed by Bio-Plex assay. The values are shown as mean ± SD from single experiments and are representative of two independent experiments. Two-way ANOVA was used for statistical analysis. Šídák’s multiple comparisons test was used for multiple comparisons between IgG control and anti-NK1.1 antibody treated mice at each time. *, *p* < 0.05; **, *p* < 0.01; ***, *p* < 0.001.

### The susceptibility of *Ifngr1*^-/-^ mice to *Ot* infection is related to bacterial strains and dose

As *Ifngr1*^-/-^ mice were highly susceptible to *Ot* infection, this mouse model may hold promise for investigating severe scrub typhus pathogenesis, bacterial virulence factors, and host immunity. To assess this potential, we compared *Ot* Karp and Gilliam (high virulent and less virulent strains), as they represent the predominant strains for human scrub typhus and are associated with different disease severity in experimental murine and non-human primate infections[18, 47]. We infected *Ifngr1*^-/-^ mice with various doses (3,000 – 30 FFU) and used WT mice as controls. As presented in **Fig. 8A**, *Ifngr1*^-/-^ mice inoculated with 3,000 FFU Karp demonstrated weight loss on days 6 or 7 p.i., and ultimately reached more than 20% body weight loss by day 12 p.i., when all mice were succumbed. *Ifngr1*^-/-^ mice inoculated with 300 or 30 FFU Karp showed similar trends of weight loss and 100% mortality on day 14 p.i.. *Ifngr1*^-/-^ mice received 3 FFU Karp showed delayed body weight loss and succumbed to the infection around day 17 p.i.. Therefore, *Ifngr1*^-/-^ mice showed clear trends of dose-dependent susceptibility to *Ot* Karp. Interestingly, *Ifngr1*^-/-^ mice inoculated with 3,000 FFU Gilliam behaved liked Karp-infected counterparts, displaying a similar trend of weight loss and animal mortality (**Fig. 8B**). *Ifngr1*^-/-^ mice received 300 FFU Gilliam demonstrated weight loss on days 6 or 7 p.i., and ultimately reached more than 20% weight loss on day 21 p.i., when all animals were died. Notably, the time window of animal death was delayed by one week in mice with 300 FFU Gilliam infection as compared to the mice with same dose of Karp infection. More notably, mice inoculated with 30 and 3 FFU Gilliam inoculation began to experience weight loss around day 15 p.i., reaching 18% and 8% weight loss on day 24 p.i., respectively; but no animal deaths were observed in these two groups by the end of the experiment (day 30 p.i.). Regarding the initial skin responses, signs of redness around injection sites were observed on days 2 or 3 p.i., regardless of bacterial strains or doses used, which disappeared after 4 days. The real skin eschar lesion was observed on day 6 p.i. after high-dose of Karp infection (3,000 FFU) as indicated by limited necrotic dermis surrounded with an indurated red halo (**Fig. 8C**). This lesion was subsequently enlarged and developed to a typical eschar that was necrotic and hardened on day 9 p.i.. This eschar lesion maintained during the infection until animal death. Mice infected with low-doses of Karp (e.g., 30 or 3 FFU) also showed eschar lesions, but lesion development was notably delayed. In contrast, 3,000 FFU Gilliam infection did not induce visible skin lesions until day 9 p.i., and the lesion scores were significantly lower than those of Karp infection (**Fig. 8D**). Gilliam infection with low-doses (300, 30, or 3 FFU) resulted in minimal or no skin lesions during days 12 to 17 p.i.; their lesion scores were also significantly lower than those of corresponding Karp infection. Together, our comparative studies with two different *Ot* strains suggested that the development of skin eschar lesions is dependent on the inoculation doses and virulence of bacterial strains.

**Fig. 8.**
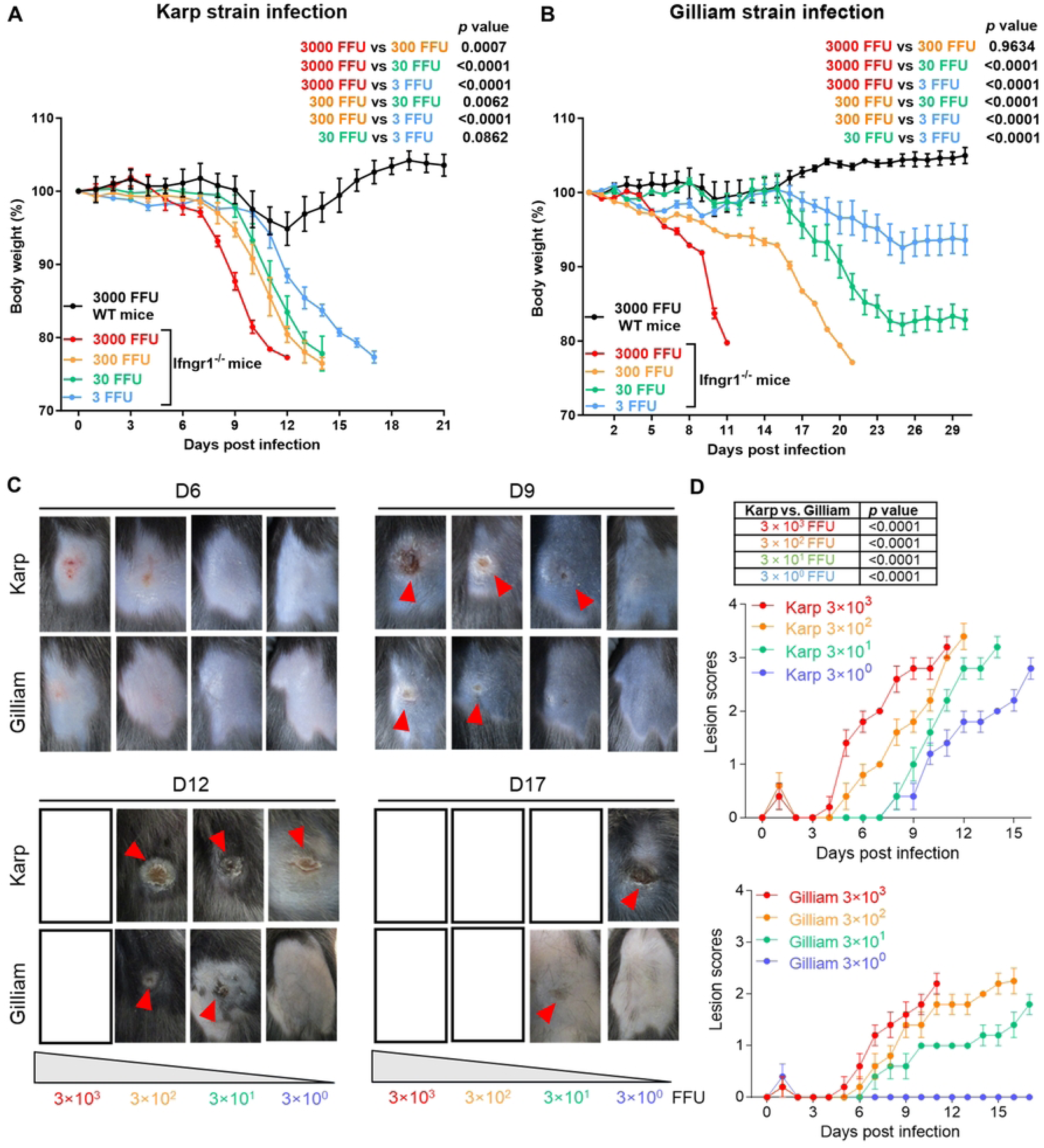
The susceptibility of *Ifngr1*^-/-^ mice to *Ot* infection is related to bacterial strains and doses. *Ifngr1*^-/-^ mice (n = 5/group) were i.d. infected with *Ot* Karp or Gilliam strain (3×10^3^, 3×10^2^, 3×10^1^ and 3×10^0^ FFU) on the flank. WT mice (n = 5/group) were i.d. infected with *Ot* Karp or Gilliam strain (3×10^3^ FFU) as controls. Body weight changes of (A) Karp and (B) Gilliam infected mice were recorded until day 30 p.i. or when all animals in the group had succumbed. (C) The photos of skin inoculation sites at days 6, 9, 12, and 17 p.i. are presented. The skin eschar-like lesions are pointed out by red arrows. The photo positions are left for blank if no mice are available at those specific time points. (D) The skin eschar-like lesion was scored for Karp and Gilliam infected *Ifngr1*^-/-^ mice (n = 5/group). The values are shown as mean ± SD from single experiments and are representative of three independent experiments. Body weight changes and skin eschar-like lesion scores were analyzed by two-way ANOVA and Šídák’s multiple comparisons test. The results of statistical analysis are presented in the graphs.

## Discussion

In the present study, we found that the lack of IFN signaling resulted in lethal *Ot* infection in an i.d. mouse model. We further demonstrated that IFN-γ, but not IFN-I, signaling was critical for host protection against *Ot* infection and skin eschar formation. The eschar lesion is the diagnostic clue in patients with acute febrile illness in areas endemic for scrub typhus; however, *Ot* infection fails to generate an eschar in immunocompetent mice[14, 16, 48]. Our study provides the first evidence for eschar formation in mice deficient in the IFN-γ-related signaling pathways, which help understand the known differences of eschar formation in human patients versus mouse models. In addition, we evaluated the severe pathology and dysregulated immune response in the absence of the IFN-γ signaling, which provides a better understanding of the detrimental vs. protective immune mechanisms in scrub typhus. We further demonstrate that NK cells act as gatekeepers in early infection and are the major source of innate IFN-γ production and contribute to bacterial control. By evaluating disease severity and eschar scores in two *Ot* strains with different virulence, we conclude that our susceptible mouse model of i.d. infection recapitulates human scrub typhus and could be beneficial for future research on *Ot* pathogenesis and host immunity.

Strong type I inflammation characterized as significantly elevated IFN-γ is a hallmark of scrub typhus in both humans and animal models[12, 22, 49, 50]. The absence of the IFN-γ signal causes severe tissue damage, uncontrolled bacterial growth, and mouse lethality (Fig. 2), indicating the crucial role of IFN-γ for host immune protection in *Ot* infection. In addition, *Ot* infection can also induce IFN-I responses via RIG-I/MAVS and cGAS/STING pathways[51]. However, the IFN-I response seems to play a redundant role in immune protection in an i.p. mouse model of *Ot* infection[51]. In agreement with that, our results demonstrate that i.d. injection of *Ot* in *Ifnar1*^-/-^ mice caused mild disease symptoms such as those observed in WT mice (Fig. 2). The dispensable role of IFN-I in vivo might be attributed to the compensatory effect of highly induced IFN-γ by *Ot*. Similar findings were also reported in an i.d. infection mouse model of *Rickettsia parkeri* (*Rp*), indicating resistance to *Rp* infection in *Ifnar1*^-/-^ mice. [29]. However, in contrast to *Ot*, *Rp* also failed to induce lethal infection in *Ifngr1*^-/-^ mice[29], indicating that IFN-I may compensate for IFN-γ in *Rp* infection, but not *Ot* infection. The reason for the discrepancies in the role of IFNs between *Ot* (Gram-negative, LPS negative) and *Rp* (Gram-negative, LPS positive) remains unclear. Since chigger-fed outbred ICR mice developed lethal scrub typhus[52], it will be important for future studies to examine whether chigger saliva components can dampen IFN-γ- or NK-mediated host defense, facilitating bacterial dissemination and replication. Further investigations into how IFNs protect hosts against *Ot* infection may reveal crucial aspects of the immune response to vector-borne diseases.

Inflammatory infiltration in major organs such as the lung and liver can cause severe tissue damage and dysregulated metabolism in *Ot* infection, leading to the high mortality rate[22]. By using i.d. inoculation and a lethal *Ifngr1*^-/-^ mouse model, we observed that neutrophils and monocytes in the lungs significantly increased at days 6 and 10 p.i. (Fig. 3), which indicates that they may play a pathogenic role and contribute to pulmonary hyperinflammation. In contrast, MФs and NK cells were reduced in the absence of the IFN-γ signaling (Fig. 3). M1 MФs, which are uniquely induced by IFN-γ plays a key role in controlling *Ot* growth[10]. NK cells are reported to be increased in scrub typhus patients and may contribute to anti-*Ot* responses via producing IFN-γ production[50]. Moreover, NK cell accumulation and activation depend on the IFN-γ signal[53], suggesting a positive feedback loop between NK cells and IFN-γ in *Ot* infection. Additionally, the number of activated T cells is comparable or even higher in *Ifngr1*^-/-^ mice as compared to WT mice in both spleen and lung. It is possible that the functionality of T cells might be dampened since we observed decreased levels of serum IL-2 (Fig. 4A), that is mainly produced by pathogen-specific T cells. Decreased serum IL-12p40 as well as IL-4, IL-5, IL-10 and TNF-α in *Ifngr1*^-/-^ mice further suggests the suboptimal T cell priming and function in the absence of IFN-γ signaling (Fig. 4A). In contrast, IL-6 and G-CSF were significantly increased in the serum of infected *Ifngr1*^-/-^ mice (Fig. 4B). IL-6 has been reported to be elevated in the lungs and liver of our i.v. mouse model[12] and may cause endothelial cell dysregulation as a key component of cytokine storm[54]. G-CSF is a growth factor that stimulates the bone marrow to produce granulocytes, such as neutrophils and release them into the bloodstream[55]. Increased G-CSF levels in the blood have been reported in a rhesus macaque model of scrub typhus[56]. Our recent study revealed that virulent Karp infection can induce higher levels of G- CSF than the less virulent Gilliam infection[18], indicating that G-CSF might be related to the hyperinflammation and immunopathology. Further investigation is warranted to reveal the unique roles of IL-6 and G-CSF and their potential as therapeutic strategies for severe scrub typhus. In addition to measuring cytokines/chemokines, we measured serum chemistry parameters in *Ot*-infected mice (Fig. 2F). The severe infection and tissue damage of *Ifngr1*^-/-^ mice were evidenced by the aberrant levels of TP, GLOB and ALB. The reduced glucose levels and ALP following infection may suggest the metabolic dysfunction and/or reduced food uptake. Unexpectedly, ALT and AMY, the liver and kidney injury markers were lower in *Ifngr1*^-/-^ mice as compared to that in WT mice. This is not consistent with the histological result which shows extensive necrosis in the absence of IFN-γ signaling (Fig. 2E) and may indicate that the upregulation of ALT and AMY might be highly dependent on IFN-γ stimulation. Dehydration may also manifest in severely infected mice, as evidenced by elevated minerals in the blood attributed to gastrointestinal dysfunction, akin to observations in patients[57, 58].

Early control of *Ot* replication and dissemination is critical for preventing the host from systemic infection and severe outcomes[18]. Innate IFN-γ production can be indued in response to intracellular bacteria by NK cells[59], playing an antimicrobial role in rickettsial infection[60]. Our in vivo data highlighted that NK cells are the main source of innate IFN-γ, playing a gatekeeper-like role in inhibiting bacterial growth or dissemination in the dLN. Furthermore, our in vitro experiment showed that IFN-γ significantly inhibited *Ot* growth in DCs (Fig. 5). Given that DCs are the key carrier mediating *Ot* dissemination[17, 38, 61], the innate IFN-γ, derived from NK cells, may control bacterial infection in the skin-dLN axis by modulating DC functions. Subsequent investigations are necessary to explore the underlying mechanism through which *Ot* activates innate immune cells (such as NK cells, NKT cells, T cells, and innate lymphoid cells) and evades the innate immune defense. In addition, the comparable bacterial burdens in the organs on days 10 and 14 p.i. suggest that adaptive T cells may be the dominant IFN-γ producers. Other myeloid cells, such as MФs may also contribute to compensatory IFN-γ production after NK cell depletion. Further investigation is warranted to understand the pathogenic vs protective role of IFN-γ in the late stage of *Ot* infection.

Skin eschar lesion in scrub typhus patients, characterized by a central black crust and peripheral erythematous rim[39], is a critical pathognomonic finding for clinical diagnosis. While eschar formation has been reported in Karp- and Gilliam-infected non-human primate models via i.d. inoculation [47, 62, 63], no eschars are observed in i.d. inoculated-immunocompetent mouse models[9, 14, 16, 17, 48, 64]. Eschar formation in *Ifngr1*^-/-^ mice highlights the important role of the IFN-γ signal in controlling skin lesions (Fig. 8), suggesting that IFN-γ levels and its downstream JAK/STAT signals as key determinants of human scrub typhus. The difference of eschar formation between humans and mice might be attributed to their distinct C-terminal tail structures of IFN-γ, which results in lower biological activity of IFN-γ in humans compared to that in mice[65, 66]. Further research is also required to identify key downstream IFN-stimulated genes in humans and mice for restricting *Ot* infection. Our detailed comparative studies with a broad range of Karp and Gilliam strains support a conclusion that the eschar formation is related to the bacterial strains. This finding is consistent with our recent report in i.v. mouse models, showing that B6 mice were susceptible to Karp, but resistant to Gilliam infection[18]. Furthermore, our finding is also supported by the clinical observation that the frequency of occurrence of eschars in scrub typhus patients might be dependent on the *Ot* strains[67]. The *Ifngr1*^-/-^ mice exhibited high susceptibility to the virulent Karp strain, with even an extremely low-dose infection (3 FFU) resulting in animal mortality. Interestingly, low-dose Gilliam infection (30 and 3 FFU) resulted in limited to invisible eschar lesions; however, mouse body weights were unable to fully recover, suggesting a potential persistent infection by Gilliam.

Additionally, our finding of a positive correlation between bacterial doses and eschar lesion scores implies that eschar might be an independent predictor of scrub typhus disease severity[68–70]. In summary, this study highlights the crucial role of IFN-γ, but not IFN-I, signaling in host protection against *Ot* infection. Our mouse models resemble skin eschar lesions and lethal infection observed in human scrub typhus. The i.d. inoculation approaches, plus comparison of different *Ot* strains and infection doses, hold promise for future immunological studies for scrub typhus, especially for the role of local and innate defense mechanisms against *Ot* infection, the bacterial virulence factors to subvert early host responses, and potential host factors for exacerbating severe scrub typhus.

## Acknowledgements

We would like to thank the UTMB Flow Cytometry and Cell Sorting Core Lab (Meredith Weglarz) for sample analyses, and Dr. David Walker for facilitating the BSL-3 research facilities. We thank Dr. Ashley Smith for assisting manuscript revision.

---

**Supplementary Fig. 1 Serum chemistry parameters in WT *and Ifngr1*^-/-^ mice following *Ot* infection.** WT *and Ifngr1*^-/-^ (n = 5/group) were i.d. infected with *Ot* Karp strain (3×10^3^ FFU) on the flank. The mouse serum chemistry profile was generated by using VetScan Comprehensive Diagnostic Profile reagent rotor. The parameters include creatinine (CRE), phosphorus (PHOS), total bilirubin (TBIL), and urea nitrogen (BUN). The values are shown as mean ± SD from single experiments and are representative of two independent experiments. Two-way ANOVA and Šídák’s multiple comparisons test were used for statistical analysis.

**Supplementary Fig. 2 Flow cytometric analysis of spleen cells.** WT and *Ifngr1*^-/-^ mice (n = 4/group) were i.d. infected with *Ot* Karp (3×10^3^ FFU). Mouse spleens were collected for preparation of single cell suspensions at days 3, 6 and 10 p.i.. Mock-infected mice were used at day 0. Cell samples were acquired by flow cytometry and data were analyzed by using Flowjo. Gating strategy is shown in Fig. 3. Dead cells and doublets were first excluded by live/dead fixable dye and FSC-H vs FSC-W/SSC-H vs SSC-W, respectively. The live and single cells were gated for activated CD4 T cells (CD3^+^CD4^+^CD44^+^CD62L^-^), activated CD8 T cells (CD3^+^CD8^+^CD44^+^CD62L^-^), and activated NK cells (CD3^-^NK1.1^+^CD69^+^). Cell numbers are presented as mean ± SD from single experiments and are representative of two independent experiments. Two-way ANOVA was used for statistical analysis. Šídák’s multiple comparisons test was used for multiple comparisons between WT B6 and *Ifngr1*^-/-^ mice at each time. ****, *p* < 0.0001.

**Supplementary Fig. 3 Bacterial burdens in WT, *Ifngr1*^-/-^ and *Ifnar1*^-/-^ mice following *Ot* infection.** WT, *Ifngr1*^-/-^ and *Ifnar1*^-/-^ mice (n = 5/group) were i.d. infected with *Ot* Karp (3×10^3^ FFU). Lung and liver tissues were harvested on day 10 p.i., followed by the measurement of bacterial burdens via qPCR. The values are shown as mean ± SD from single experiments and are representative of two independent experiments. A one-way ANOVA statistical analysis with a Tukey’s multiple comparisons test was used for multiple comparisons. ****, *p* < 0.0001; ns, no significance.

**Supplementary Fig. 4 NK cells depletion by anti-NK1.1 antibody.** B6 mice (n = 5-6/group) were i.p. treated with either IgG control or anti-NK1.1 neutralizing antibody (200 µg/mouse) on days -3 and -1 and were euthanized on day 0. Single-cell suspensions were prepared by using the dLN, lung and spleen, followed by flow cytometric analysis. The representative images of flow cytometry were shown. The cell percentages are shown as mean ± SD. Two-tails student t-test was used for comparisons between IgG control and anti-NK1.1 antibody treated samples. **, *p* < 0.01; ****, *p* < 0.0001.

**Supplementary Fig. 5 Bio-Plex assay of anti-NK1.1 treated mice.** B6 mice (5-6/group) were i.d. infected with *Ot* Karp (3×10^3^ FFU) and were i.p. treated with either IgG control or anti-NK1.1 neutralizing antibody (200 µg/mouse/treatment) every other day starting from 3 day prior to infection. Mice were euthanized at days 3, 6, 10, and 14 p.i. for tissue/blood harvest. Mouse serum was assessed for by Bio-Plex assay. The values are shown as mean ± SD from single experiments and are representative of two independent experiments. Two-way ANOVA was used for statistical analysis. Šídák’s multiple comparisons test was used for multiple comparisons between IgG control and anti-NK1.1 antibody treated mice at each time.

## References

1. Luce-Fedrow A, Lehman ML, Kelly DJ, Mullins K, Maina AN, Stewart RL, et al. A Review of Scrub Typhus (*Orientia tsutsugamushi* and Related Organisms): Then, Now, and Tomorrow. Trop Med Infect Dis. 2018;3(1):8.

2. Chakraborty S, Sarma N. Scrub Typhus: An Emerging Threat. Indian J Dermatol. 2017;62(5):478–85.

3. Varghese GM, Trowbridge P, Janardhanan J, Thomas K, Peter JV, Mathews P, et al. Clinical profile and improving mortality trend of scrub typhus in South India. Int J Infect Dis. 2014;23:39–43.

4. Taylor AJ, Paris DH, Newton PN. A Systematic Review of Mortality from Untreated Scrub Typhus (*Orientia tsutsugamushi*). PLoS Negl Trop Dis. 2015;9(8):e0003971.

5. Jiang J, Richards AL. Scrub Typhus: No Longer Restricted to the Tsutsugamushi Triangle. Trop Med Infect Dis. 2018;3(1):11.

6. Weitzel T, Dittrich S, Lopez J, Phuklia W, Martinez-Valdebenito C, Velasquez K, et al. Endemic Scrub Typhus in South America. N Engl J Med. 2016;375(10):954–61.

7. Chen K, Travanty NV, Garshong R, Crossley D, Wasserberg G, Apperson CS, et al. Detection of *Orientia* spp. Bacteria in Field-Collected Free-Living *Eutrombicula* Chigger Mites, United States. Emerg Infect Dis. 2023;29(8):1676–9.

8. Paris DH, Shelite TR, Day NP, Walker DH. Unresolved problems related to scrub typhus: a seriously neglected life-threatening disease. Am J Trop Med Hyg. 2013;89(2):301–7.

9. Mendell NL, Xu G, Shelite TR, Bouyer DH, Walker DH. A Murine Model of Waning Scrub Typhus Cross-Protection between Heterologous Strains of *Orientia tsutsugamushi*. Pathogens. 2022;11(5):512.

10. Trent B, Liang Y, Xing Y, Esqueda M, Wei Y, Cho NH, et al. Polarized lung inflammation and Tie2/angiopoietin-mediated endothelial dysfunction during severe *Orientia tsutsugamushi* infection. PLoS Negl Trop Dis. 2020;14(3):e0007675.

11. Soong L, Shelite TR, Xing Y, Kodakandla H, Liang Y, Trent BJ, et al. Type 1-skewed neuroinflammation and vascular damage associated with *Orientia tsutsugamushi* infection in mice. PLoS Negl Trop Dis. 2017;11(7):e0005765.

12. Soong L, Wang H, Shelite TR, Liang Y, Mendell NL, Sun J, et al. Strong type 1, but impaired type 2, immune responses contribute to *Orientia tsutsugamushi*-induced pathology in mice. PLoS Negl Trop Dis. 2014;8(9):e3191.

13. Shelite TR, Saito TB, Mendell NL, Gong B, Xu G, Soong L, et al. Hematogenously disseminated *Orientia tsutsugamushi*-infected murine model of scrub typhus [corrected]. PLoS Negl Trop Dis. 2014;8(7):e2966.

14. Keller CA, Hauptmann M, Kolbaum J, Gharaibeh M, Neumann M, Glatzel M, et al. Dissemination of *Orientia tsutsugamushi* and inflammatory responses in a murine model of scrub typhus. PLoS Negl Trop Dis. 2014;8(8):e3064.

15. Sunyakumthorn P, Paris DH, Chan TC, Jones M, Luce-Fedrow A, Chattopadhyay S, et al. An intradermal inoculation model of scrub typhus in Swiss CD-1 mice demonstrates more rapid dissemination of virulent strains of *Orientia tsutsugamushi*. PLoS One. 2013;8(1):e54570.

16. Soong L, Mendell NL, Olano JP, Rockx-Brouwer D, Xu G, Goez-Rivillas Y, et al. An Intradermal Inoculation Mouse Model for Immunological Investigations of Acute Scrub Typhus and Persistent Infection. PLoS Negl Trop Dis. 2016;10(8):e0004884.

17. Liang Y, Wang H, Gonzales C, Thiriot J, Sunyakumthorn P, Melby PC, et al. CCR7/dendritic cell axis mediates early bacterial dissemination in *Orientia tsutsugamushi*-infected mice. Front Immunol. 2022;13:1061031.

18. Thiriot JD, Liang Y, Gonzales C, Sun J, Yu X, Soong L. Differential cellular immune responses against *Orientia tsutsugamushi* Karp and Gilliam strains following acute infection in mice. PLoS Negl Trop Dis. 2023;17(12):e0011445.

19. Xu G, Mendell NL, Liang Y, Shelite TR, Goez-Rivillas Y, Soong L, et al. CD8+ T cells provide immune protection against murine disseminated endotheliotropic *Orientia tsutsugamushi* infection. PLoS Negl Trop Dis. 2017;11(7):e0005763.

20. Hauptmann M, Kolbaum J, Lilla S, Wozniak D, Gharaibeh M, Fleischer B, et al. Protective and Pathogenic Roles of CD8+ T Lymphocytes in Murine *Orientia tsutsugamushi* Infection. PLoS Negl Trop Dis. 2016;10(9):e0004991.

21. Shelite TR, Liang Y, Wang H, Mendell NL, Trent BJ, Sun J, et al. IL-33-Dependent Endothelial Activation Contributes to Apoptosis and Renal Injury in *Orientia tsutsugamushi*-Infected Mice. PLoS Negl Trop Dis. 2016;10(3):e0004467.

22. Soong L. Dysregulated Th1 Immune and Vascular Responses in Scrub Typhus Pathogenesis. J Immunol. 2018;200(4):1233–40.

23. Tantibhedhyangkul W, Ben Amara A, Textoris J, Gorvel L, Ghigo E, Capo C, et al. *Orientia tsutsugamushi*, the causative agent of scrub typhus, induces an inflammatory program in human macrophages. Microb Pathog. 2013;55:55–63.

24. Tantibhedhyangkul W, Prachason T, Waywa D, El Filali A, Ghigo E, Thongnoppakhun W, et al. *Orientia tsutsugamushi* stimulates an original gene expression program in monocytes: relationship with gene expression in patients with scrub typhus. PLoS Negl Trop Dis. 2011;5(5):e1028.

25. Thiriot J, Liang Y, Fisher J, Walker DH, Soong L. Host transcriptomic profiling of CD-1 outbred mice with severe clinical outcomes following infection with *Orientia tsutsugamushi*. PLoS Negl Trop Dis. 2022;16(11):e0010459.

26. Fisher J, Card G, Liang Y, Trent B, Rosenzweig H, Soong L. *Orientia tsutsugamushi* selectively stimulates the C-type lectin receptor Mincle and type 1-skewed proinflammatory immune responses. PLoS Pathog. 2021;17(7):e1009782.

27. Liang Y, Aditi, Onyoni F, Wang H, Gonzales C, Sunyakumthorn P, et al. Brain transcriptomics reveal the activation of neuroinflammation pathways during acute *Orientia tsutsugamushi* infection in mice. Front Immunol. 2023;14:1194881.

28. Fisher J, Gonzales C, Chroust Z, Liang Y, Soong L. *Orientia tsutsugamushi* Infection Stimulates Syk-Dependent Responses and Innate Cytosolic Defenses in Macrophages. Pathogens. 2022;12(1):53.

29. Burke TP, Engstrom P, Tran CJ, Langohr IM, Glasner DR, Espinosa DA, et al. Interferon receptor-deficient mice are susceptible to eschar-associated rickettsiosis. Elife. 2021;10:e67029.

30. Burke TP, Engstrom P, Chavez RA, Fonbuena JA, Vance RE, Welch MD. Inflammasome-mediated antagonism of type I interferon enhances *Rickettsia* pathogenesis. Nat Microbiol. 2020;5(5):688–96.

31. Li H, Jerrells TR, Spitalny GL, Walker DH. Gamma interferon as a crucial host defense against Rickettsia conorii in vivo. Infect Immun. 1987;55(5):1252–5.

32. Liang Y, Fisher J, Gonzales C, Trent B, Card G, Sun J, et al. Distinct Role of TNFR1 and TNFR2 in Protective Immunity Against *Orientia tsutsugamushi* Infection in Mice. Front Immunol. 2022;13:867924.

33. Wang X, Spandidos A, Wang H, Seed B. PrimerBank: a PCR primer database for quantitative gene expression analysis, 2012 update. Nucleic Acids Res. 2012;40(Database issue):D1144–9.

34. Jie Z, Liang Y, Yi P, Tang H, Soong L, Cong Y, et al. Retinoic Acid Regulates Immune Responses by Promoting IL-22 and Modulating S100 Proteins in Viral Hepatitis. J Immunol. 2017;198(9):3448–60.

35. Liu Z, Gu Y, Shin A, Zhang S, Ginhoux F. Analysis of Myeloid Cells in Mouse Tissues with Flow Cytometry. STAR Protoc. 2020;1(1):100029.

36. Trent B, Fisher J, Soong L. Scrub Typhus Pathogenesis: Innate Immune Response and Lung Injury During *Orientia tsutsugamushi* Infection. Front Microbiol. 2019;10:2065.

37. Kim DM, Won KJ, Park CY, Yu KD, Kim HS, Yang TY, et al. Distribution of eschars on the body of scrub typhus patients: a prospective study. Am J Trop Med Hyg. 2007;76(5):806–9.

38. Paris DH, Phetsouvanh R, Tanganuchitcharnchai A, Jones M, Jenjaroen K, Vongsouvath M, et al. *Orientia tsutsugamushi* in human scrub typhus eschars shows tropism for dendritic cells and monocytes rather than endothelium. PLoS Negl Trop Dis. 2012;6(1):e1466.

39. Park J, Woo SH, Lee CS. Evolution of Eschar in Scrub Typhus. Am J Trop Med Hyg. 2016;95(6):1223–4.

40. Platanias LC. Mechanisms of type-I- and type-II-interferon-mediated signalling. Nat Rev Immunol. 2005;5(5):375–86.

41. Jiang L, Morris EK, Aguilera-Olvera R, Zhang Z, Chan TC, Shashikumar S, et al. Dissemination of *Orientia tsutsugamushi*, a Causative Agent of Scrub Typhus, and Immunological Responses in the Humanized DRAGA Mouse. Front Immunol. 2018;9:816.

42. Knipfer L, Schulz-Kuhnt A, Kindermann M, Greif V, Symowski C, Voehringer D, et al. A CCL1/CCR8-dependent feed-forward mechanism drives ILC2 functions in type 2-mediated inflammation. J Exp Med. 2019;216(12):2763–77.

43. Barsheshet Y, Wildbaum G, Levy E, Vitenshtein A, Akinseye C, Griggs J, et al. CCR8(+)FOXp3(+) T(reg) cells as master drivers of immune regulation. Proc Natl Acad Sci U S A. 2017;114(23):6086–91.

44. Piao HL, Tao Y, Zhu R, Wang SC, Tang CL, Fu Q, et al. The CXCL12/CXCR4 axis is involved in the maintenance of Th2 bias at the maternal/fetal interface in early human pregnancy. Cell Mol Immunol. 2012;9(5):423–30.

45. Yoon HJ, Lee MS, Ki M, Ihm C, Kim D, Kim Y, et al. Does IL-17 play a role in hepatic dysfunction of scrub typhus patients? Vector Borne Zoonotic Dis. 2010;10(3):231–5.

46. Iwasaki H, Mizoguchi J, Takada N, Tai K, Ikegaya S, Ueda T. Correlation between the concentrations of tumor necrosis factor-alpha and the severity of disease in patients infected with *Orientia tsutsugamushi*. Int J Infect Dis. 2010;14(4):e328–33.

47. Inthawong M, Sunyakumthorn P, Wongwairot S, Anantatat T, Dunachie SJ, Im-Erbsin R, et al. A time-course comparative clinical and immune response evaluation study between the human pathogenic *Orientia tsutsugamushi* strains: Karp and Gilliam in a rhesus macaque (Macaca mulatta) model. PLoS Negl Trop Dis. 2022;16(8):e0010611.

48. Mendell NL, Bouyer DH, Walker DH. Murine models of scrub typhus associated with host control of *Orientia tsutsugamushi* infection. PLoS Negl Trop Dis. 2017;11(3):e0005453.

49. Munch CC, Upadhaya BP, Rayamajhee B, Adhikari A, Munch M, En-Nosse N, et al. Multiple *Orientia* clusters and Th1-skewed chemokine profile: a cross-sectional study in patients with scrub typhus from Nepal. Int J Infect Dis. 2023;128:78–87.

50. Kang SJ, Jin HM, Cho YN, Kim SE, Kim UJ, Park KH, et al. Increased level and interferon-gamma production of circulating natural killer cells in patients with scrub typhus. PLoS Negl Trop Dis. 2017;11(7):e0005815.

51. Min CK, Kim HI, Ha NY, Kim Y, Kwon EK, Yen NTH, et al. A Type I Interferon and IL-10 Induced by *Orientia tsutsugamushi* Infection Suppresses Antigen-Specific T Cells and Their Memory Responses. Front Immunol. 2018;9:2022.

52. Lurchachaiwong W, Monkanna T, Leepitakrat S, Ponlawat A, Sattabongkot J, Schuster AL, et al. Variable clinical responses of a scrub typhus outbred mouse model to feeding by *Orientia tsutsugamushi* infected mites. Exp Appl Acarol. 2012;58(1):23–34.

53. Lin Q, Rong L, Jia X, Li R, Yu B, Hu J, et al. IFN-gamma-dependent NK cell activation is essential to metastasis suppression by engineered *Salmonella*. Nat Commun. 2021;12(1):2537.

54. Kang S, Kishimoto T. Interplay between interleukin-6 signaling and the vascular endothelium in cytokine storms. Exp Mol Med. 2021;53(7):1116–23.

55. Bendall LJ, Bradstock KF. G-CSF: From granulopoietic stimulant to bone marrow stem cell mobilizing agent. Cytokine Growth Factor Rev. 2014;25(4):355–67.

56. Linsuwanon P, Wongwairot S, Auysawasdi N, Monkanna T, Richards AL, Leepitakrat S, et al. Establishment of a Rhesus Macaque Model for Scrub Typhus Transmission: Pilot Study to Evaluate the Minimal *Orientia tsutsugamushi* Transmission Time by Leptotrombidium chiangraiensis Chiggers. Pathogens. 2021;10(8):1028.

57. Aung T, Supanaranond W, Phumiratanaprapin W, Phonrat B, Chinprasatsak S, Ratanajaratroj N. Gastrointestinal manifestations of septic patients with scrub typhus in Maharat Nakhon Ratchasima Hospital. Southeast Asian J Trop Med Public Health. 2004;35(4):845–51.

58. Kim DM, Yun NR, Lim SC. Neuritis and gastrointestinal hemorrhage in scrub typhus patients. Am J Trop Med Hyg. 2015;92(1):145–7.

59. Thale C, Kiderlen AF. Sources of interferon-gamma (IFN-gamma) in early immune response to *Listeria monocytogenes*. Immunobiology. 2005;210(9):673–83.

60. Fang R, Ismail N, Walker DH. Contribution of NK cells to the innate phase of host protection against an intracellular bacterium targeting systemic endothelium. Am J Pathol. 2012;181(1):185–95.

61. Choi JH, Cheong TC, Ha NY, Ko Y, Cho CH, Jeon JH, et al. *Orientia tsutsugamushi* subverts dendritic cell functions by escaping from autophagy and impairing their migration. PLoS Negl Trop Dis. 2013;7(1):e1981.

62. Walsh DS, Delacruz EC, Abalos RM, Tan EV, Jiang J, Richards AL, et al. Clinical and histological features of inoculation site skin lesions in cynomolgus monkeys experimentally infected with *Orientia tsutsugamushi*. Vector Borne Zoonotic Dis. 2007;7(4):547–54.

63. Paris DH, Chattopadhyay S, Jiang J, Nawtaisong P, Lee JS, Tan E, et al. A nonhuman primate scrub typhus model: protective immune responses induced by pKarp47 DNA vaccination in cynomolgus macaques. J Immunol. 2015;194(4):1702–16.

64. Luce-Fedrow A, Chattopadhyay S, Chan TC, Pearson G, Patton JB, Richards AL. Comparison of Lethal and Nonlethal Mouse Models of *Orientia tsutsugamushi* Infection Reveals T-Cell Population-Associated Cytokine Signatures Correlated with Lethality and Protection. Trop Med Infect Dis. 2021;6(3):121.

65. Lundell D, Lunn C, Dalgarno D, Fossetta J, Greenberg R, Reim R, et al. The carboxyl-terminal region of human interferon gamma is important for biological activity: mutagenic and NMR analysis. Protein Eng. 1991;4(3):335–41.

66. Zhao W, Valencia AZ, Melby PC. Biological activity of hamster interferon-gamma is modulated by the carboxyl-terminal tail. Cytokine. 2006;34(5-6):243–51.

67. Kim DM, Yun NR, Neupane GP, Shin SH, Ryu SY, Yoon HJ, et al. Differences in clinical features according to Boryoung and Karp genotypes of *Orientia tsutsugamushi*. PLoS One. 2011;6(8):e22731.

68. Lee CS, Hwang JH, Lee HB, Kwon KS. Risk factors leading to fatal outcome in scrub typhus patients. Am J Trop Med Hyg. 2009;81(3):484–8.

69. Chauhan V, Thakur A, Thakur S. Eschar is associated with poor prognosis in scrub typhus. Indian J Med Res. 2017;145(5):693–6.

70. Kim DM, Kim SW, Choi SH, Yun NR. Clinical and laboratory findings associated with severe scrub typhus. BMC Infect Dis. 2010;10:108.

